# High-throughput discovery of inhibitory protein fragments with AlphaFold

**DOI:** 10.1101/2023.12.19.572389

**Authors:** Andrew Savinov, Sebastian Swanson, Amy E. Keating, Gene-Wei Li

**Affiliations:** Department of Biology, Massachusetts Institute of Technology, Cambridge, MA, USA; Department of Biological Engineering, Massachusetts Institute of Technology, Cambridge, MA, USA; Koch Center for Integrative Cancer Research, Massachusetts Institute of Technology, Cambridge, MA, USA

**Keywords:** Inhibitory peptides, protein interactions, deep mutational scanning, massively parallel, FragFold

## Abstract

Peptides can bind to specific sites on larger proteins and thereby function as inhibitors and regulatory elements. Peptide fragments of larger proteins are particularly attractive for achieving these functions due to their inherent potential to form native-like binding interactions. Recently developed experimental approaches allow for high-throughput measurement of protein fragment inhibitory activity in living cells. However, it has thus far not been possible to predict *de novo* which of the many possible protein fragments bind to protein targets, let alone act as inhibitors. We have developed a computational method, FragFold, that employs AlphaFold to predict protein fragment binding to full-length proteins in a high-throughput manner. Applying FragFold to thousands of fragments tiling across diverse proteins revealed peaks of predicted binding along each protein sequence. Comparisons with experimental measurements establish that our approach is a sensitive predictor of fragment function: Evaluating inhibitory fragments from known protein-protein interaction interfaces, we find 87% are predicted by FragFold to bind in a native-like mode. Across full protein sequences, 68% of FragFold-predicted binding peaks match experimentally measured inhibitory peaks. Deep mutational scanning experiments support the predicted binding modes and uncover superior inhibitory peptides in high throughput. Further, FragFold is able to predict previously unknown protein binding modes, explaining prior genetic and biochemical data. The success rate of FragFold demonstrates that this computational approach should be broadly applicable for discovering inhibitory protein fragments across proteomes.

**Significance Statement:** Peptides can regulate protein interactions by binding to specific interfaces, and fragments of larger proteins have high potential to function in this manner. Recently developed experimental methods allow massively parallel measurement of protein fragment-based inhibition *in vivo*. However, we have lacked comparable computational methods to predict which protein fragments act as inhibitors and how they bind. Here we report a new approach, FragFold, which leverages high-throughput AlphaFold predictions of protein – fragment binding to tackle these problems at scale. FragFold is successful at predicting inhibitory protein fragments and their binding modes across diverse protein structures and functions. This new approach stands to enable proteome-wide discovery of inhibitory protein fragments and aid the interpretation of high-throughput experimental measurements of inhibitory activity.

**Classification:** Biological Sciences / Biophysics and Computational Biology

## Introduction

Peptides have great potential for modulating function in living cells by binding to and regulating target proteins. Indeed, organisms have evolved a large array of polypeptides of ≤100 amino acids, known variously as biological peptides, small proteins, miniproteins, or microproteins (1–4). Evolved miniproteins can be as structurally simple as single alpha helices (5–7) and often control protein function in response to stress conditions (5–7). Synthetic random peptides have also been shown to produce diverse biological activities (11–13), and recently, it has become feasible to design miniproteins and functional peptides, with many exciting developments in this space (14–16).

Peptides that can emulate, modulate, and rewire native protein-protein interactions – and the general features that permit such activities – are therefore of great interest. One class of peptides with an inherent potential for this kind of functionality are protein fragments derived from full-length sequences. Massively parallel measurements of fragment-based growth inhibition *in vivo* allow a systematic interrogation of such peptides (17–20). This approach revealed that protein fragments can act as dominant negative inhibitors. Savinov *et al.* demonstrated that these fragments are able to titrate native interactions of their parental proteins in living cells (20). This work also uncovered other principles of fragment-based inhibition, including the role of protein fragment length (20). Notably, generic biophysical properties of protein fragments (such as charge, hydrophobicity, and secondary structure) explained very little of the fragment-to-fragment variation in inhibitory activity. These observations suggest that specific features of fragments and their bespoke binding interfaces largely determine which fragments are potent inhibitors.

A computational approach to predicting protein fragment binding would be a powerful tool to discover inhibitory fragments across proteomes and provide structural models for inhibitory interactions. Such an approach would complement high-throughput measurements of peptide function. Experimental measurements would provide the ground truth, allowing development of a model, and computational prediction would expand the reach of experimental methods by pre-identifying likely inhibitory regions of proteins for screening. Predicting binding using AlphaFold (21) is attractive, as this model was trained using native structural data (22) and coevolutionary sequence information (e.g., ref. (23)) to extract features of natural proteins. AlphaFold has already shown promise for predicting binding modes for peptides engaging proteins (24–27), importantly including peptides not present in the training set (24, 27), and has been fruitfully applied to the prediction of protein-protein interactions more broadly (28).

Savinov *et al.* demonstrated an association between AlphaFold-predicted native-like protein-peptide binding and measured inhibitory activity for 6 protein fragments of *Escherichia coli* FtsZ, a structural protein essential for cell division (20). Here we build on that work by applying high-throughput AlphaFold computational predictions to 2,277 tiling protein fragments covering the full sequences of six diverse proteins, mirroring the massively parallel experimental measurements (20). Analysis of the results reveals that computational predictions of protein-fragment binding are in striking agreement with the *in vivo* inhibitory activities of protein fragments that are derived from protein-protein binding interfaces, with the inhibitory fragments predicted to bind in a native-like manner. We also leverage our approach to predict novel binding modes of fragments from the intrinsically disordered C-terminal tail of FtsZ to several protein binding partners. Excitingly, these new structural models explain biological activity of the FtsZ C-terminus (and its peptide fragments) consistent with genetic and biochemical evidence, providing a demonstration of the generalizability of our approach to protein interactions of unknown structure. We additionally perform deep mutational scanning experiments on several inhibitory protein fragments, showing that the spectrum of mutational effects is consistent with FragFold-predicted binding modes and discover many mutations that yield more potent inhibitory peptides.

Overall, we demonstrate the power of AlphaFold to predict protein fragments that inhibit protein interactions in living cells and the complementarity of this computational approach with high-throughput experimental measurements. These results pave the way for proteome-wide discovery of inhibitory protein fragments and provide a promising path to uncover the underlying molecular features of potent peptide inhibitors.

## Results

### An approach for high-throughput AlphaFold prediction of peptide-protein interactions

We have lacked scalable computational approaches to predict which protein fragments are inhibitory and to generate models for their binding modes. We tackled this problem by developing a highly parallelized approach leveraging the ColabFold implementation of AlphaFold2 (21, 29). In all cases, we used the AlphaFold2 monomer model weights, which were derived from training exclusively on single polypeptide chain data (21, 24) (in contrast with AlphaFold-Multimer weights that were determined using protein complex structures (30)), to ensure that native binding interactions were not memorized during model training. Protein-peptide binding predictions in AlphaFold2 were run using the established approach implemented in ColabFold, in which multiple polypeptides are modeled explicitly as separate chains (29) (*Materials and Methods*).

To run AlphaFold2 (hereafter, AlphaFold) on many protein fragments tiling across proteins of interest, we modified the slow multiple-sequence alignment (MSA) generation step. Taking advantage of the fact that the fragments are segments of full-length proteins, we pre-generated full-protein MSAs and then pruned them to generate each fragment MSA, before running AlphaFold. We fed each precalculated fragment MSA and target protein MSA into a fragment + target protein AlphaFold prediction (**Fig. 1**, *Materials and Methods*). The target protein was in some cases the parent protein of the fragment (for homomeric interactions) and in other cases one of its interaction partners (for heteromeric interactions), typically based on known interaction partners from protein-protein interaction data. In contrast to previous approaches that evaluated binding predictions primarily based on AlphaFold confidence metrics (24, 25, 31), we focused on properties of the structural models AlphaFold produced, particularly the number of binding contacts between the protein fragment and protein target (*N*_contacts_); we further weighted this number by the interface-predicted template modeling (ipTM) score, a confidence metric for multichain structural models (30). We used weighted *N*_contacts_ to generate plots of predicted binding as a function of tiling fragment sequence position (**Fig. 1**). We termed this approach FragFold. We compared FragFold predictions to *in vivo* inhibitory effects of the same set of protein fragments from massively parallel measurements (**Fig. 1**, ref. (20)).

**Figure 1.**
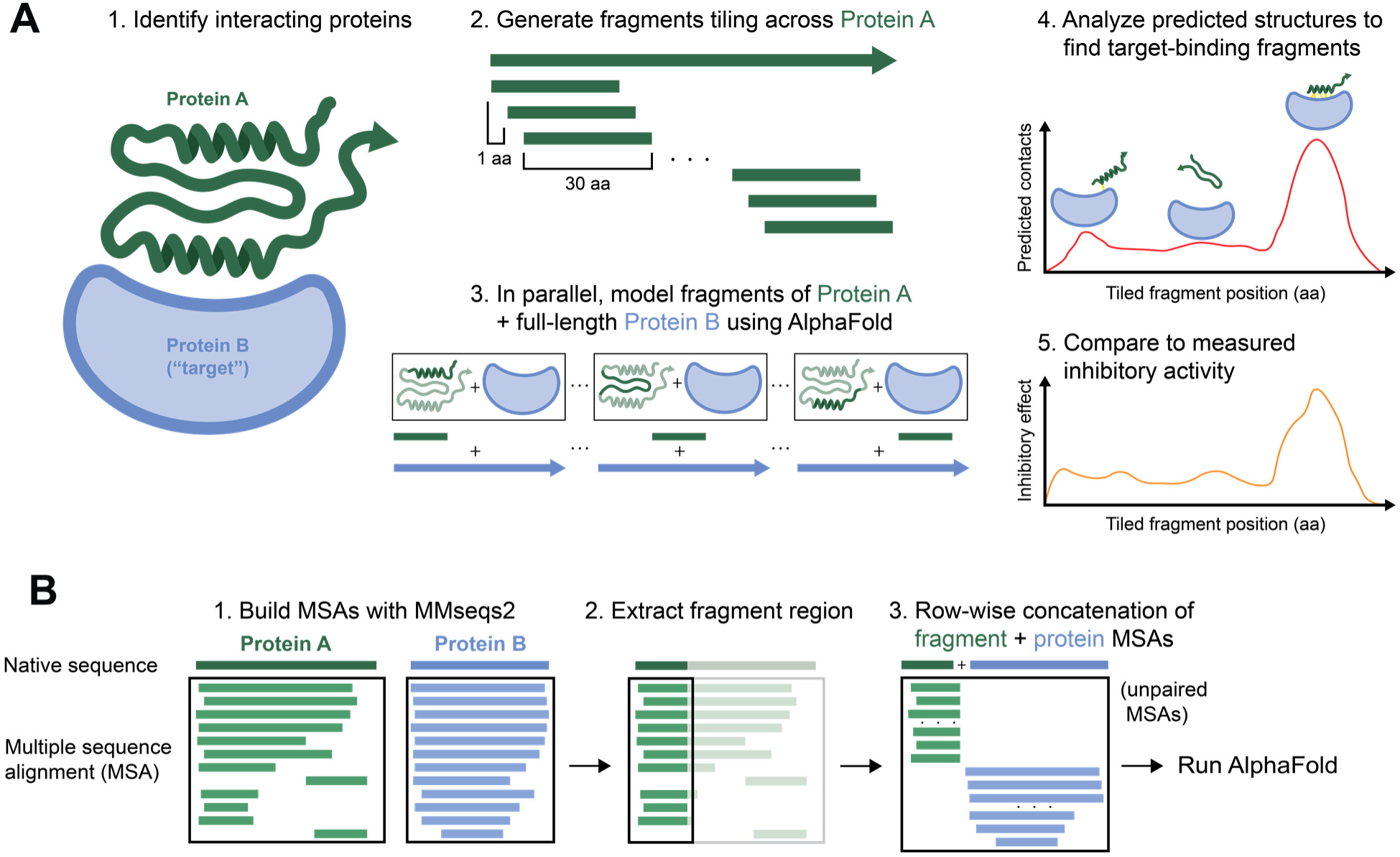
FragFold enables high-throughput prediction of protein fragment binding to protein targets. (**A**) Schematic of the FragFold pipeline. Protein pairs of interest known or suspected to interact are selected and fragmented *in silico*. Each tiling fragment is modeled together with the target interaction partner using AlphaFold2 (see *Materials and Methods*). Predicted structures are analyzed based on a combination of binding contacts and AlphaFold confidence metrics. The results can be compared to experimental measurements of protein fragment inhibitory activity in cells. (**B**) Schematic of the MSA-building scheme employed in FragFold to overcome the computational bottleneck of MSA generation for each fragment + protein pair.

### Computational prediction of peptide-protein binding recapitulates inhibitory fragment peaks across the FtsZ protein sequence

We first applied FragFold to predict protein fragment binding in the context of experimentally mapped inhibitory fragments that originate from structurally resolved protein-protein interaction interfaces. Such complexes provide an ideal test case where the relationship between binding and inhibition is most straightforward. We initially focused on the cell division protein FtsZ because fragments from across the filament polymerization interface act as inhibitors *in vivo* (20). Savinov *et al.* previously found that tiling inhibitory protein fragments form visually evident peaks around maxima in the inhibitory activity; in the case of FtsZ these peaks identified 4 distinct regions of the filament interface (**1**, **1′**, **2**, and **2′**; ref (20)). We thus applied FragFold to perform a computational scan for fragments of FtsZ binding to full-length FtsZ, examining all possible 30-amino acid (aa) fragments with a 1-aa offset, as in the experiments (20). For each peak of inhibitory activity across the polymerization interface, there was a corresponding observed peak (local maximum) in weighted *N*_contacts_ calculated for tiling protein fragments (**Fig. 2*A***). Unweighted *N*_contacts_ predicted the same inhibitory peaks, whereas the AlphaFold confidence metrics ipTM (30) and pLDDT (28) alone did not (**Fig. S1**).

**Figure 2.**
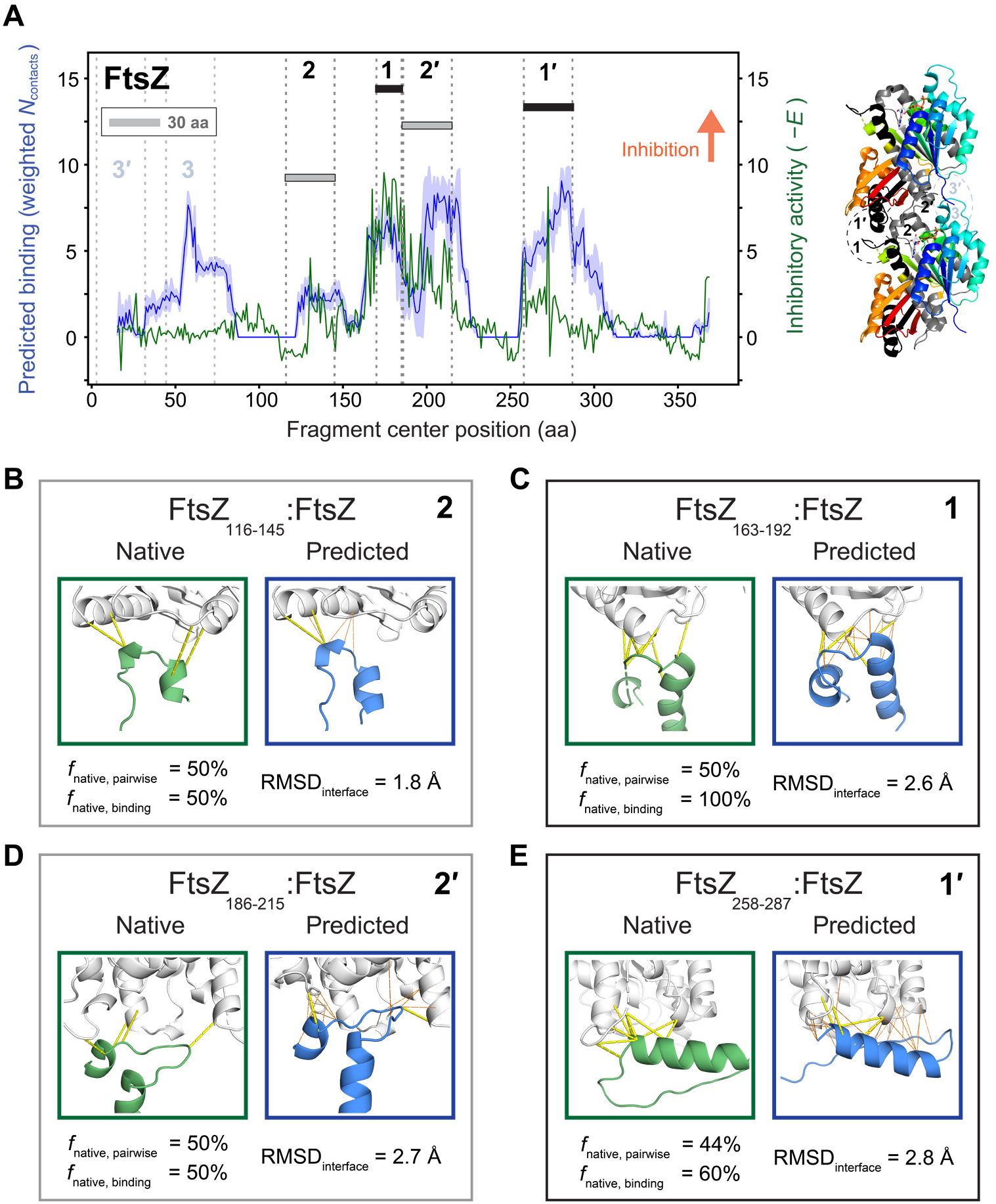
FragFold predicts corresponding binding peaks for all measured inhibitory peaks across the FtsZ filament interface, with native-like binding modes. (**A**) *Left*: Predicted target protein binding from FragFold (blue curves) vs. experimentally measured *in vivo* inhibitory activity (green curves, from ref. 20) for 30-aa fragment scans across *E. coli* FtsZ. Darker line and lighter outline for predicted binding data indicate mean and 95% C.I., respectively, across the 5 top-ranked structural models generated by AlphaFold. Filament interface regions **2**, **1**, **1′**, and **2′** are indicated. *Right*: crystallographic structure of two adjacent monomers of the FtsZ filament (PDB ID 6unx, ref. 32). Protein is colored from blue to red from the N- to the C-terminus. Regions corresponding to interface regions are annotated and highlighted in the same black/grey color scheme as in the fragment scan plot (*Left*). aa, amino acids. (**B**) – (**E**) Native, experimentally determined structures from the full FtsZ-FtsZ complex (green outlines; PDB ID 6unx) vs. FragFold-predicted structures of FtsZ fragment + full-length FtsZ (blue outlines) for protein fragments corresponding to each of the inhibitory peaks **2**, **1**, **1′**, and **2′**. Metrics for the extent of native-like binding are shown below. For each inhibitory peak (**B** – **E**), the representative measured inhibitory fragment is indicated, and the corresponding predicted binding fragment from the same structural region is as listed in **Table 1**. Pairwise native residue contacts are shown as dashed yellow lines; additional predicted native binding site contacts are shown as thinner dashed orange lines.

**Table 1.** AlphaFold predicts native-like binding for the majority of inhibitory fragment peaks that correspond to known protein-protein interaction interfaces.

| Fragmented protein | Binding partner | Measured inhibitory fragment (aa) | Predicted fragment (aa) | $f_{\text{native, pairwise}}$ | $f_{\text{native, binding}}$ | RMSD<sub>interface</sub> (Å) | $N_{\text{contacts}}$ | Weighted $N_{\text{contacts}}$ |
| --- | --- | --- | --- | --- | --- | --- | --- | --- |
| **FtsZ** | FtsZ | 116 - 145 | 118 - 147 | 50% | **50%** | 1.7 | 6 | **2.2** |
| **FtsZ** | FtsZ | 163 - 192 | 162 - 191 | 50% | **100%** | 2.6 | 18 | **8.4** |
| **FtsZ** | FtsZ | 186 - 215 | 186 - 215 | 50% | **50%** | 2.7 | 16 | **7.0** |
| **FtsZ** | FtsZ | 258 - 287 | 271 - 300 | 44% | **60%** | 2.8 | 27 | **10.2** |
| **RplL** | RplL | 5 - 34 | 1 - 30 | 61% | **93%** | 1.2 | 24 | **5.8** |
| **RplL** | RplJ | 5 - 34 | 5 - 34 | 0% | **67%** | 4.6 | 14 | **2.1** |
| **GyrA** | GyrB | 12 - 41 | 12 - 41 | 47% | **89%** | 3.0 | 39 | **16.1** |
| **GyrA** | GyrA | 371 - 400 | 371 - 400 | 100% | **100%** | 0.8 | 7 | **2.6** |
| **LptG** | LptF | 317 - 330 | 319 - 332 | 60% | **75%** | 2.37 | 12 | **2.9** |
| **Ssb** | Ssb | 52 - 81 | 53 - 82 | 100% | **100%** | 2.3 | 5 | **3.1** |
| **GroEL** | GroEL | 3 - 32 | 3 - 32 | 88% | **75%** | 0.4 | 7 | **3.4** |
| **GroEL** | GroEL | 486 - 515 | 493 - 522 | 75% | **75%** | 0.5 | 24 | **12.7** |
| **GroEL** | GroEL | 87 - 116 | 87 - 116 | 0% | 0% | 0.7 | 3 | 1.7 |
| **GroEL** | GroES | 229 - 258 | 226 - 255 | 11% | 17% | 3.2 | 1 | 0 |
| **GroEL** | GroES | 246 - 275 | 246 - 275 | 38% | **100%** | 3.2 | 16 | **5.1** |

For each experimental inhibitory peak identified as mapping to a PPI, the best matching fragment from FragFold and its native-like binding characteristics (as well as predicted $N_{\text{contacts}}$ and weighted $N_{\text{contacts}}$ ) are listed. Bolded values indicate predictions that meet the criteria for binding (weighted $N_{\text{contacts}} > 2$ ) and native-like interaction mode ( $f_{\text{native, binding}} \geq 50\%$ ).

Having determined the concordance of predicted binding and measured inhibitory activity for these interface regions, we next analyzed the predicted binding modes of fragments from each inhibitory peak. Strikingly, the FragFold predictions for sites **1**, **1′**, **2**, and **2′** placed the peptide in a geometry similar to the native binding mode seen in a crystallographic structure of the filament (32) (**Fig. 2*B*–*E***). To quantify this similarity, we calculated the fraction of native pairwise contacts between fragment and target residues recapitulated in the FragFold binding models, *f*_native, pairwise_ (with native pairwise contacts defined as those seen in experimental structures of the full-length protein-protein interaction). As another metric of similarity, we determined the fraction of target protein residues from the native binding site predicted to be bound by any residue of the fragment, *f*_native, binding_. Finally, we additionally calculated the root mean-square deviation (RMSD) of the FragFold-predicted fragment-target interface compared to the native structure, RMSD_interface_ (**Fig. 2*B*–*E***; *Materials and Methods*). All four major peaks yielded predictions with RMSD_interface_ < 3 Å, *f*_native, binding_ ≥ 50%, and *f*_native, pairwise_ > 40%. We also observed a predicted binding peak centered around FtsZ residue 60 with no corresponding inhibitory peak (**Fig. 2*A***). This region maps to a loop involved in filament formation (interaction site **3**) and was predicted to bind in a native-like geometry by FragFold. However, this region has high B-factors in the experimental structure, suggesting it contributes little to the binding energy (20, 32). Thus, this example represents a case where a plausible predicted binder does not inhibit the native protein-protein interaction.

### FragFold predicts protein-protein interaction inhibitory fragments across diverse proteins

We next sought to determine how generally FragFold can predict inhibitory peaks mapping to protein interaction interfaces, and their binding modes, across structurally and functionally diverse proteins. We therefore extended our computational approach to fragment scans of several additional *E. coli* proteins: 50S ribosomal subunit protein L7/L12 (RplL), DNA gyrase subunit A (GyrA), single-stranded DNA binding protein (Ssb), lipopolysaccharide transport protein subunit G (LptG), and the GroEL/ES chaperone subunit GroEL. We previously obtained experimental measurements of *in vivo* inhibitory activity for fragments spanning each of these targets (20), revealing 36 inhibitory fragment peaks. Each of these proteins forms structurally characterized protein-protein interactions (PPIs), which allowed us to compare predicted binding modes to native structural data for complexes.

Across these diverse proteins, we evaluated predicted binding for all inhibitory peaks that we could assign to known PPIs based on experimental structures (14 inhibitory peaks corresponding to 15 PPIs; ref. (20), **Fig. 3**, **Table 1**; green highlighted regions in **Fig. S2**). Overall, we found that ∼87% (13/15) of PPI-inhibitory fragment-protein interactions were predicted to bind in a native-like geometry, with weighted *N*_contacts_ > 2 and *f*_native, binding_ ≥ 50%) (**Fig. 3**, **Table 1**).

**Figure 3.**
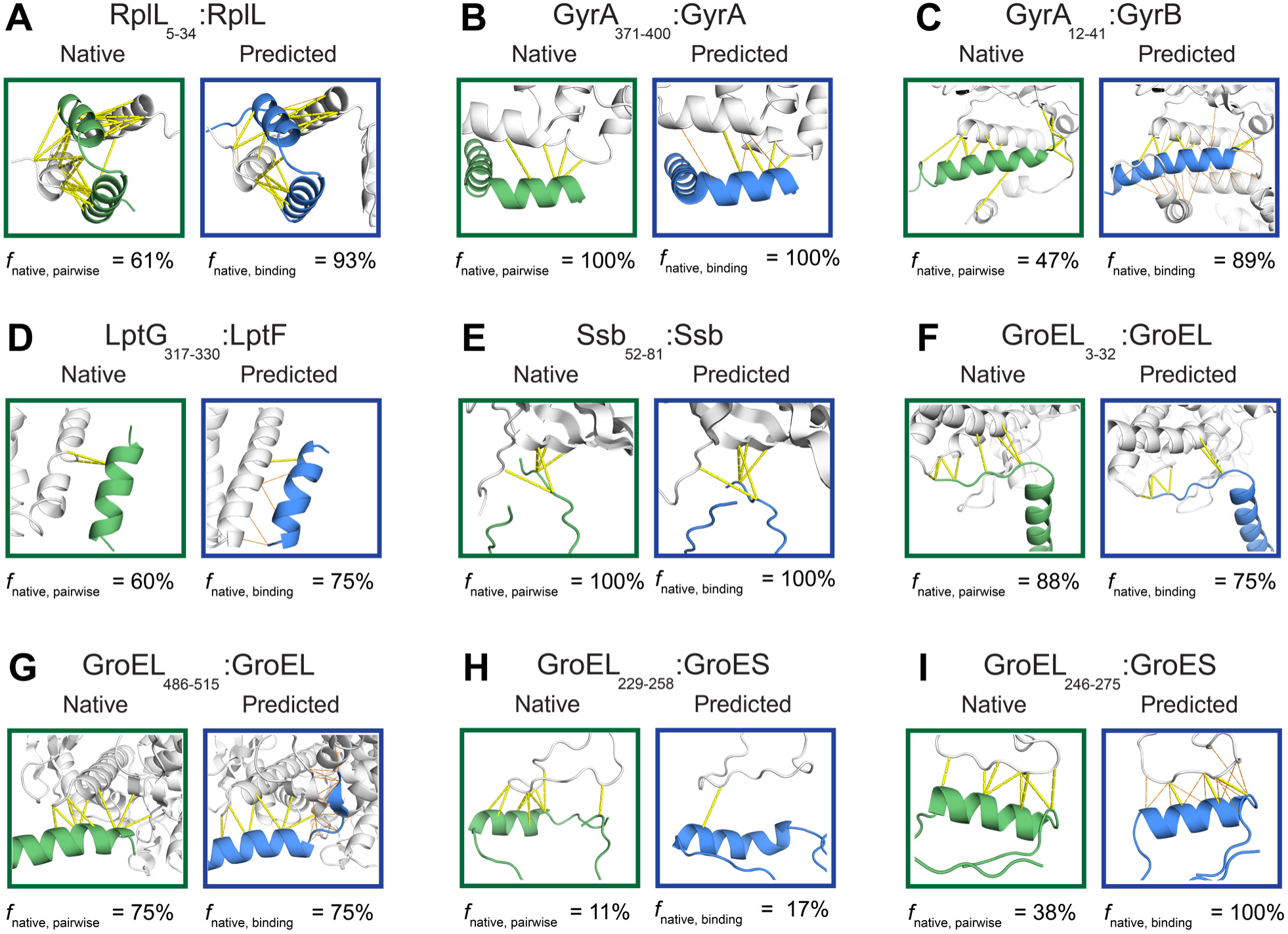
FragFold successfully predicts native-like protein interaction inhibitors across structurally and functionally diverse proteins. (**A**) – (**I**) Native, experimentally determined structures from full-length protein complexes (green outlines) vs. FragFold-predicted structures of protein fragments + full-length target protein (blue outlines) for protein fragments corresponding to each of the indicated inhibitory peaks identified in *in vivo* experiments (ref. 20). Metrics for extent of native-like binding are shown below each structural comparison. For each inhibitory peak (**A** – **I**), the representative measured inhibitory fragment is indicated, and the corresponding predicted binding fragment from the same structural region is as listed in **Table 1**. Pairwise native residue contacts are shown as dashed yellow lines; additional predicted native binding site contacts are shown as thinner dashed orange lines.

For instance, for RplL, we saw that the single inhibitory peak corresponding to the RplL dimerization domain was predicted to bind to RplL in a native-like manner, with RMSD_interface_ = 1.2 Å and *f*_native, binding_ = 93% (**Fig. 3*A***). The binding of 50S ribosomal subunit L10 (RplJ) to this same fragment of RplL was predicted as well, though with orientational deviation from the native mode (RMSD_interface_ = 4.6 Å and *f*_native, binding_ = 67%; **Table 1**). For GyrA, AlphaFold predicted native-like binding to GyrB for the inhibitory fragment peak at residues 12-41 (*f*_native, binding_ = 89%) and native-like binding to GyrA at the C-gate dimerization site (experimental inhibitory peak at residues 371-400; *f*_native, binding_ = 100%) (**Fig. 3*B*,*C***). For LptG, our computational predictions successfully yielded the single inhibitory peak corresponding to a protein-protein interaction with LptF (LptG residues 317-330; *f*_native, binding_ = 75%) (**Fig. 3*D***). For Ssb, the lone inhibitory peak was likewise reproduced computationally (residues 52-81; *f*_native, binding_ = 100%) (**Fig. 3*E***, **Table 1**). Finally, for GroEL, fragments corresponding to inhibitory peaks at residues 3-32 and 486-515 mapping to GroEL-GroEL polymerization interactions were predicted to bind in native-like modes (each with *f*_native, binding_ = 75%; **Fig. 3*F*,*G***) but GroEL_87-116_: GroEL was not (*f*_native, binding_ = 0%) (**Table 1**). The GroEL-GroES interface inhibitory peak fragments (GroEL residues 229-258, 246-275) were predicted also, with *f*_native, binding_ = 100% for fragment 246-275, but *f*_native, binding_ of only 17% for fragment 229-258. Despite this, the interface RMSD was quite reasonable (3.2 Å) for the 229-258 prediction (**Fig. 3*H*,*I***). Interface RMSD was also low for the GroEL_87-116_: GroEL prediction (0.7 Å), indicating that in both of these cases where the prediction failed using the weighted *N*_contacts_ and *f*_native, binding_ criteria, AlphaFold nonetheless came close to predicting a native-like binding mode.

### Automated prediction of inhibitory fragments across diverse proteins suggests generalizability across proteomes

The results from the structural predictions of inhibitory peaks mapping to known protein-protein interaction interfaces showed that FragFold can discern these types of inhibitory peaks with a high success rate (**Table 1**). However, not all inhibitory peaks in the experimental data had obvious structural interpretations based on the protein complex structures examined, and more generally the set of known protein interaction structures is incomplete. Furthermore, the relationship between (predicted) protein binding and inhibitory function is not necessarily straightforward, with binding at known interaction interfaces representing the simplest case. We therefore sought to develop an automated approach that could be applied to discover inhibitory protein fragments from computational fragment scans of arbitrary proteins, given candidate interaction partners.

To do so, we began from our computational results alone and automatically determined predicted binding peaks as clusters of overlapping protein fragments predicted to bind the target protein with a sufficient number of contacts at a common binding site. Computationally predicted target-binding fragment clusters were then compared to experimentally determined inhibitory fragment peaks. To make this process as unsupervised as possible, we developed an algorithm to automatedly determine experimental inhibitory peaks from massively parallel fragment scans, which was able to reproduce all 14 previously manually assigned inhibitory peaks across 6 proteins (*Materials and Methods*; **Table 1** and ref. (20)). In addition to these 14 peaks, 22 additional visually evident inhibitory peaks not mapping to known PPIs were predicted by FragFold (**Fig. S2**, green lines). Comparison of the predicted binding clusters with the experimental inhibitory peaks revealed that all previously identified PPI-inhibitory peaks (**Table 1**) were predicted by FragFold using this automated pipeline (**Fig. S2**, green highlighted regions).

We sought to estimate how often a FragFold-predicted protein-binding fragment is inhibitory using an automated approach. We based this assessment purely on the overlap of predicted binding clusters and ground-truth experimental inhibitory peaks across all investigated protein sequences. We optimized the tradeoff between prediction sensitivity (fraction of all inhibitory fragment peaks mapping to a corresponding predicted binding cluster, *f* _inhibitory peaks predicted_) and specificity (fraction of all predicted binding clusters (peaks) that map to inhibitory peaks, *f* _predicted binding clusters inhibitory_) by adjusting the peak calling criteria (**Fig. S3**; *Materials and Methods*). Using our chosen parameters (including at least 6 fragments per cluster, *N*_contacts_ ≥ 3, and weighted *N*_contacts_ ≥ 3; *Materials and Methods*), we found that *f* _inhibitory peaks predicted_ was ∼64% (23/36) (**Table 2**, **Fig. S2**). This included all protein interaction-inhibitory peaks in **Table 1**. At this level of sensitivity, predictions also showed good specificity: we found that *f* _predicted binding clusters inhibitory_ was ∼68% (23/34) (**Table 2**, **Fig. S2**). As a further measure of specificity, we found that *f* _predicted binding clusters inhibitory_ (68% over all proteins) was higher than expected from random placement of peaks across all proteins investigated (51%) (**Table S1**). Evaluating the ability of FragFold to predict 36 peaks across all 6 proteins, this difference from the null model was statistically significant (*p* = 0.048; *Materials and Methods*). Similarly, across all proteins, *f* _inhibitory peaks predicted_ (64%) was also significantly larger than expected from random placement (49%; *p* = 0.035). Together, these results suggest that AlphaFold can successfully predict many protein fragment inhibitory peaks across diverse proteins.

**Table 2.** FragFold systematically predicts inhibitory protein fragment peaks.

| Fragmented protein | $f_{\text{inhibitory peaks predicted}}$ | $f_{\text{predicted binding clusters inhibitory}}$ |
| --- | --- | --- |
| All proteins | 64% | 68% |
| FtsZ | 86% | 71% |
| GroEL | 44% | 67% |
| GyrA | 54% | 70% |
| LptG | 75% | 57% |
| RplL | 100% | 100% |
| Ssb | 100% | 67% |

The fraction of measured inhibitory peaks that matched corresponding predicted binding clusters ( $f_{\text{inhibitory peaks predicted}}$ ) as well as the fraction of predicted binding clusters (peaks) that matched corresponding experimentally determined inhibitory peaks ( $f_{\text{predicted binding clusters inhibitory}}$ ) are listed. These results are presented for all fragmented proteins ("All proteins"), as well as for each protein individually.

### Specificity of FragFold predictions

We tested the specificity of FragFold predictions by performing fragment scans against noncognate protein partners (**Fig. S4**). FtsZ fragments tiling the full protein sequence were not predicted to interact with enhanced GFP (eGFP); likewise, fragments of eGFP were not predicted to bind to FtsZ (**Fig. S4*A*–*B***). Repeating this test for GyrA, no GyrA fragments were predicted to bind to FtsZ (**Fig. S4*C***). When scanning FtsZ fragments for binding to GyrA, some binding peaks were predicted, but these peaks were weaker than the specific peaks and not aligned in sequence position with the experimentally measured inhibitory peaks (**Fig. S4*D***). We note as well that these noncognate predicted interaction peaks were uniformly narrow compared to typical native interaction-mapped peaks. This result suggests that a greater number of overlapping fragments predicted to bind a target is indicative of a *bona fide*, or at any rate stronger, protein-peptide interaction. The structural features of these predicted nonspecific fragment binding modes include parallel alpha-helix and beta-sheet interactions (**Fig. S4*E*–*H***), suggesting that they are plausible modes of weak nonspecific association. The predicted binding peaks of eGFP fragments to eGFP (**Fig. S4*B***, grey) likely reflect the known propensity of GFP to dimerize, with the peaks centered around residues 157 and 191 (**Fig. S4*B*,*I*–*J***) including key residues involved in dimerization (33, 34).

We performed additional FragFold predictions of binding of all possible 30-aa fragments FtsZ, GyrA, RplL, and eGFP to 7-8 “off-target” protein partners each (**Fig. S5**), and leveraged our automated peak-calling approaches to further interrogate the specificity of FragFold using these results. We calculated a normalized predicted peak density, ρ_peaks_ = *N*_peaks_ / *N*_fragments_ / *L*_targets(aa)_ (where *L*_targets(aa)_ is the total length of the target protein sequences), with dimensions of peaks / fragment / aa, for both on-target (ρ_peaks,on-target_) and off-target (ρ_peaks,off-target_) predictions (**Table S2**). We then determined ρ_peaks,on-target_ / ρ_peaks,off-target_ as a metric for the specificity of predictions for each of FtsZ, GyrA, and RplL. We found that on-target predictions were 10-fold more prevalent than off-target predictions for FtsZ, 12-fold for GyrA, and 4-fold for RplL. Further, we note that the only significant peaks called for eGFP came from the GFP-GFP dimerization interactions discussed above. This systematic investigation reinforces that FragFold is generally specific in its predictions of inhibitory protein fragments. However, the level of specificity varies from protein to protein.

### AlphaFold predicts previously unknown binding modes of the intrinsically disordered C-terminal tail of FtsZ in agreement with orthogonal functional measurements

The success of FragFold in predicting inhibitory fragment peaks and native-like binding modes across diverse proteins – despite training only on folding of single-chain polypeptides (21, 24) – prompted us to wonder whether AlphaFold can successfully predict previously unknown fragment binding modes. Success in this task would further support generalizability of FragFold for predicting likely inhibitory protein fragment binders of any protein of interest. To investigate this question, we focused on the intrinsically disordered C-terminal tail of FtsZ, comprised of residues 317-383 (35, 36). This region of the protein is known to be important for regulatory interactions with other divisome proteins including FtsA, MinC, and ZipA in *E. coli* (36), but the intrinsically disordered tail is completely unresolved in FtsZ crystal structures (32). Extensive prior genetic and biochemical data on the interaction modes of the C-terminal tail provide a point of comparison for FragFold-predicted binding models, making this example an ideal test case for FragFold prediction of novel interactions. Previously, Savinov *et al.* identified a peak of *in vivo* inhibitory activity corresponding to fragments from the extreme C-terminus of FtsZ (residues 354-383; **Fig. 2**, ref. (20)). Investigating the predicted binding of tiling FtsZ peptide fragments to FtsZ itself, FtsA, MinC, and ZipA, we found that there were clear binding peaks to FtsZ and FtsA mapping to this C-terminal inhibitory peak; there was also a weak signature for predicted binding to ZipA in the same region (**Fig. 4*A***).

**Figure 4.**
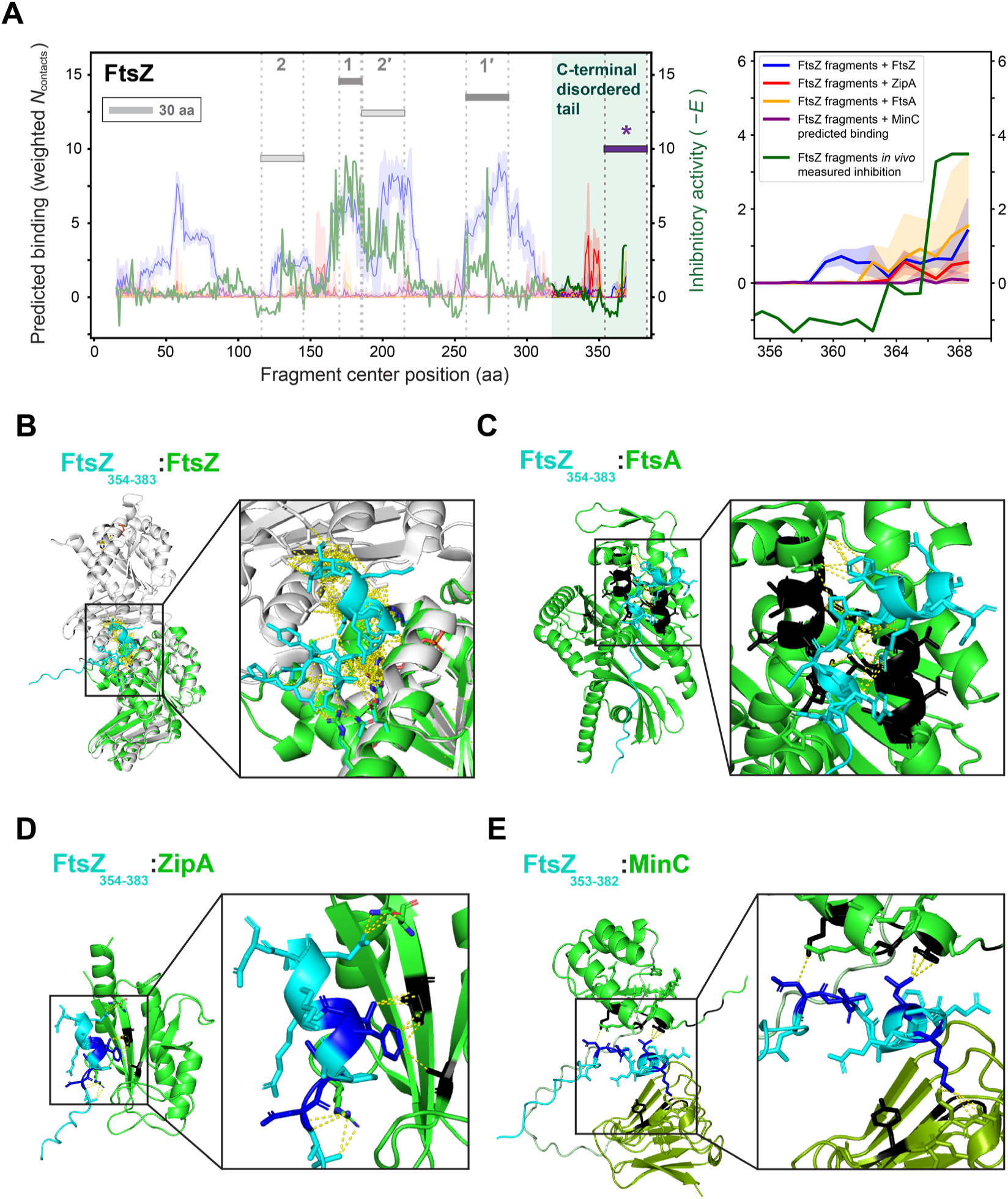
FragFold predicts novel binding modes for the intrinsically disordered FtsZ tail in agreement with genetic and biochemical evidence. (**A**) *Left*: Predicted target protein binding from FragFold (blue, red, tan, and purple curves) vs. experimentally measured *in vivo* inhibitory activity (green curves, from ref. 20) for 30 aa fragment scans across *E. coli* FtsZ with partner proteins FtsZ, ZipA, FtsA, and MinC as indicated in the legend on the *Right*. Darker line and lighter outline for predicted binding data indicate mean and 95% C.I., respectively, across the 5 top-ranked structural models generated by AlphaFold. Filament formation interface regions **2**, **1**, **1′**, and **2′** are indicated. The highlighted region of the plot corresponds to the intrinsically disordered C-terminal tail. The indicated region **\*** (purple) corresponds to the peak of fragment inhibitory activity in the C-terminal tail. *Right*: Inset from the plot on the *Left* focusing on the C-terminal inhibitory peak. aa, amino acids. (**B**) – (**E**) AlphaFold-predicted structures of indicated C-terminal FtsZ fragments (cyan) bound to various divisome proteins (green). Contacts between residue pairs are shown as dotted yellow lines. (**B**) Grey: FtsZ dimer structure from PDB ID 6unx (32), shown overlaid with AlphaFold model. (**C**) – (**E**) Black: regions of FtsA, ZipA, or MinC previously identified to play a role in FtsZ interactions (see main text); (**D**) – (**E**) Blue: regions of FtsZ previously shown to play a role in interactions with ZipA or MinC, respectively (main text). (**E**) MinC^N^ domain shown in bright green, MinC^C^ domain in olive green; interdomain linker in pale green.

We analyzed the predicted structural features of C-terminal FtsZ fragment interactions with divisome proteins. Considering first the predicted binding mode to full-length FtsZ, we found that C-terminal fragments are predicted to bind to the GTPase active site at interaction site **2**-**2′**, with fragments predicted to both occlude the GTPase binding site and prevent binding to the T7 loop of the adjacent FtsZ monomer in the filament (**Fig. 4*B***). Both features of this binding mode are expected to prevent filament polymerization and hence are consistent with the observed inhibitory activity. Excitingly, this novel structural model agrees with biochemical measurements indicating that the C-terminal tail plays an autoregulatory role in FtsZ filament assembly and acts as an auto-inhibitor of GTPase activity (37, 38). In addition, the binding mode predicted by AlphaFold is consistent with molecular dynamics simulations indicating that the C-terminal end of the C-terminal tail binds in the vicinity of the GTPase active site (38), with our results providing the first detailed molecular model for this interaction. Notably, predictions of full-length FtsZ folding and dimerization fail to predict this C-terminal tail binding mode (**Fig. S6**), demonstrating that AlphaFold predictions using short protein fragments can reveal interaction interfaces invisible when using full protein sequences.

In the case of FtsA, AlphaFold predicts FtsZ C-terminal fragment binding to subdomain 2B in the vicinity of FtsA residues 234-245 and 294-304 (**Fig. 4*C***). This new molecular model for FtsZ-FtsA binding agrees strikingly well with the results of genetic screens for FtsA mutants disrupting the FtsA-FtsZ interaction (39), which outlined a binding pocket in the same region of the 2B subdomain (**Fig. 4*C***, black). Notably, the predicted binding mode we report here places the FtsZ C-terminal peptide near the ATP-binding site of FtsA, potentially inhibiting nucleotide binding. ATP binding by membrane-associated FtsA is important for FtsA membrane remodeling and Z-ring constriction (40), suggesting that this structural model for the FtsZ tail – FtsA binding mode reflects a key regulatory interaction between these two cell division proteins.

We also considered the predicted binding of C-terminal FtsZ fragments to ZipA (**Fig. 4*D***). In this case an experimental structural model is available (41). Although the predicted binding peak is weak, the predicted binding mode broadly agrees with the experimental structure, with the C-terminal tail peptide of FtsZ binding to the same beta-sheet face of ZipA, but in a different orientation. Furthermore, two of the three most important FtsZ residues for ZipA binding revealed in previous work (Phe377 and Leu378) are also key binding residues in our structure; as in the crystallographic structure, they make contacts with Ala246 and Phe269 of ZipA (**Fig. 4*D***, black and blue residues). Interestingly, Mosyak and colleagues previously found that ZipA binds C-terminal FtsZ fragments weakly (K_D_ ∼20-35 µM, ref. (41)), and the crystallographic peptide-protein interface contains substantial water content, with an elevated B-factor for the FtsZ fragment. Together, these features suggest that various orientations of peptide binding may be plausible at this weak interface, and the structural model produced by AlphaFold represents a reasonable binding mode. We noted as well the predicted ZipA-binding peaks for FtsZ fragments centered around residues 343 and 347 (**Fig. 4*A***). Interestingly, these correspond to predicted binding to the same ZipA interface as fragment 354-383 (**Fig. S7)**. This suggests ZipA may act as an anchor for two different regions of the FtsZ intrinsically disordered tail.

Finally, we investigated binding between the FtsZ C-terminus and MinC. Across the top 5 AlphaFold models, the average binding peak was very weak. But a local maximum in weighted *N*_contacts_ (**Fig. 4*A***, *right*) prompted us to investigate further. The highest-ranked model corresponds to the C-terminal FtsZ fragment 352-383 making 3 residue-residue contacts with MinC (**Fig. 4*E***). The structural model is intriguing, with the FtsZ tail binding in between the two domains of MinC (MinC^N^ and MinC^C^). The predicted binding mode is in good agreement with genetic and biochemical analyses of FtsZ and MinC (42–44). All three key binding residues predicted for FtsZ (D373, V378, and M380; **Fig. 4*E***, blue), as well as their partners on MinC^N^ (K35 and F41; **Fig. 4*E***, black), were previously shown to be critical for this protein-protein interaction. Although the exact MinC^C^ residues bound by FtsZ in our model (L203 and D205) were not previously shown to be involved in this interaction, a nearby residue on the same face of MinC^C^ (Y201) was, suggesting only a small adjustment to this binding mode would be needed to explain the role of this residue as well. We note that the AlphaFold model employs some of the residues shown to have a role in FtsZ binding but not others, which is also consistent with prior measurements: in addition to binding the intrinsically disordered C-terminal tail, MinC also binds elsewhere on FtsZ as part of its function in filament disassembly (43).

Our results for the example of the FtsZ C-terminal tail demonstrate that AlphaFold can predict novel peptide-protein interactions that inhibit key protein functions in cells. The agreement of these novel structural models with orthogonal experimental results suggests that they capture key features of the interactions in question.

### FragFold-predicted binding modes of protein fragments are comparable those of known peptide binders

We next evaluated the relative binding strength and physical plausibility of FragFold predictions by estimating their binding energy using Rosetta. Compared to 87 protein-peptide complexes with known experimental structures (from the PepPro database (45)), predicted protein fragment binding modes exhibit a similar energy distribution (dG_separated/dSASAx100), with binding energies of -2.9 ± 0.8 dG_separated/dSASAx100 (mean ± s.d.) for FragFold-modeled protein fragments and -3.6 ± 0.8 for the PepPro peptides (amounting to a ∼20% difference) and considerable overlap in the energy distributions (**Fig. 5**). This analysis supports the FragFold-predicted binding modes as physically and biochemically reasonable. The slightly weaker binding energy is consistent with the fact that protein fragments have not evolved for optimal binding in isolation, as well as possible imperfections in predicted binding modes compared to experimental structures of bound peptides. This gap is further reduced when the predicted structural models are allowed to relax under the influence of the Rosetta energy function (with default parameters, *Materials and Methods*), allowing for tweaking of the AlphaFold-predicted binding modes: this procedure yields dG_separated/ dSASAx100 of -3.3 ± 0.7 (mean ± s.d.) for FragFold-modeled protein fragments and -3.8 ± 0.7 for the PepPro peptides (**Fig. 5**). This relaxation amounted to typically small interface rearrangements (1.2 ± 3.0 Å, median ± s.d.; falling to 1.0 ± 0.6 Å when two outliers with RMSD > 3.0 Å are removed) (**Fig. S8**). Indeed, details including sidechain orientations and hydrogen bonding patterns at interfaces are similar between FragFold models and FragFold models with subsequent Rosetta relaxation (**Fig. S8**). The small rearrangements seen with default Rosetta relaxation parameters suggest that fine-tuning of the binding modes generated by AlphaFold directly can be productive, but is not necessary to obtain a reasonable model for protein fragment binding.

**Figure 5.**
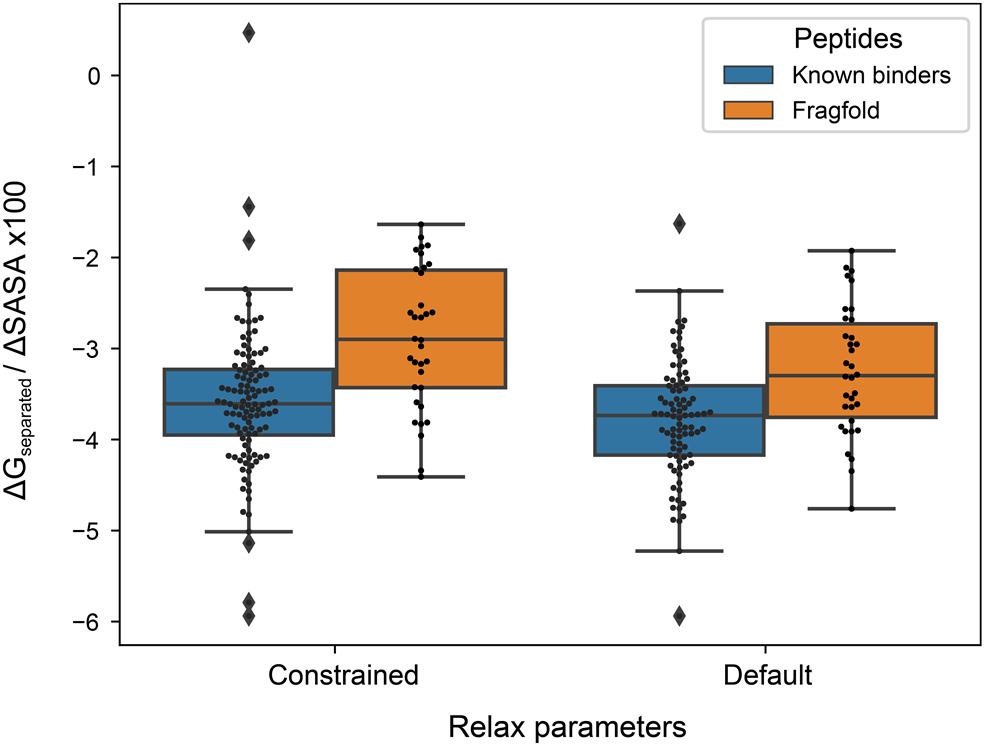
FragFold binding models exhibit comparable Rosetta binding energies to known peptide binding modes. Rosetta-modeled binding energy (dG_separated/dSASAx100) compared for FragFold structures (orange) and PepPro database peptide complexes (blue), with model geometry either tightly constrained (*Left*) or allowed to relax with standard parameters (*Right*).

### Comparison to Peptiderive predictions

We were curious how the results of our FragFold predictions, as well as the original experimental data on protein fragment inhibitory activity *in vivo* (20), compared to an alternative approach for predicting putative peptide fragment binders: the Peptiderive protocol implemented in Rosetta (46). Peptiderive uses a structure-based approach, identifying protein-protein interfaces in a given structure to predict interface-binding peptide fragments, and scoring them using the Rosetta energy function (47). Overall, we found that FragFold recapitulates all binding fragments predicted by Peptiderive, and additionally identifies several others that Peptiderive fails to predict (**Fig. S9**). Specifically, Peptiderive predicts 18/23 (78%) of the experimental inhibitory peaks predicted by FragFold (corresponding to 50% of experimental inhibitory peaks in total). We note as well that the massively parallel experimental data on in-cell fragment inhibitory activity (20) provides, to our knowledge, the first large-scale test of function of protein fragments predicted by Peptiderive. It is exciting to see that this method can also systematically identify *in vivo* fragment inhibitors. The concordance between high-throughput experimental measurements, FragFold, and orthogonal predictions from Peptiderive reinforces the finding that small protein fragments are pervasively functional as inhibitors of native interactions in living cells (20).

### Deep mutational scanning of inhibitory protein fragments reveals functional roles of each residue *in vivo* and supports FragFold-predicted fragment binding modes

To experimentally investigate the determinants of binding and inhibition by protein fragments, we performed a deep mutational scan of fragments spanning several FragFold-predicted inhibitory peaks across diverse proteins. This scan included 8 inhibitory peaks mapping to experimentally determined structural interfaces from five proteins, as well as the C-terminal inhibitory peak of FtsZ mapping to novel interaction modes determined solely by FragFold. Each residue of each selected fragment was mutated to 4-7 possible alternative amino acids with varied physicochemical properties (including an alanine scan, positive and negative charges, bulky, and aromatic residues at each position; *Materials and Methods*). Mutant and wild-type (wt) fragments were encoded on an expression vector, and a high-throughput assay of *in vivo* inhibitory effects was performed (20). For each residue of each fragment tested, the mutational sensitivity Δ_Inhibition_ was then calculated as the change in inhibitory effect (growth assay enrichment) due to the mutation, compared to the wt reference (i.e., Δ_Inhibition_ = *E*_mutant_ – *E*_wt_, where *E* is the enrichment). We thus determined mutational sensitivity at amino-acid resolution, revealing how mutations to each position of an inhibitory fragment can alter its function in living cells.

Overlaying the average mutational sensitivity for each residue across the fragment-protein complex structures revealed a rich landscape of functional effects (**Fig. 6**, **Fig. S10**). Overall, 66% of all residues exhibited substantial sensitivity with average effects of ⟨Δ_Inhibition_⟩ > 0.5 (representing mutant fragments with *reduced* inhibitory activity) or ⟨Δ_Inhibition_⟩ < –0.5 (mutants with *increased* inhibitory activity). Positions with changes tending to produce loss of function were slightly more common (36.4%) compared to residues where mutations on average increase inhibitory activity (29.4%), but both were prevalent across fragment sequences. The high frequency of sites that are mutationally sensitive may suggest a high degree of cooperativity in folding and binding (e.g., coupled folding and binding) of these short protein fragments.

**Figure 6.**
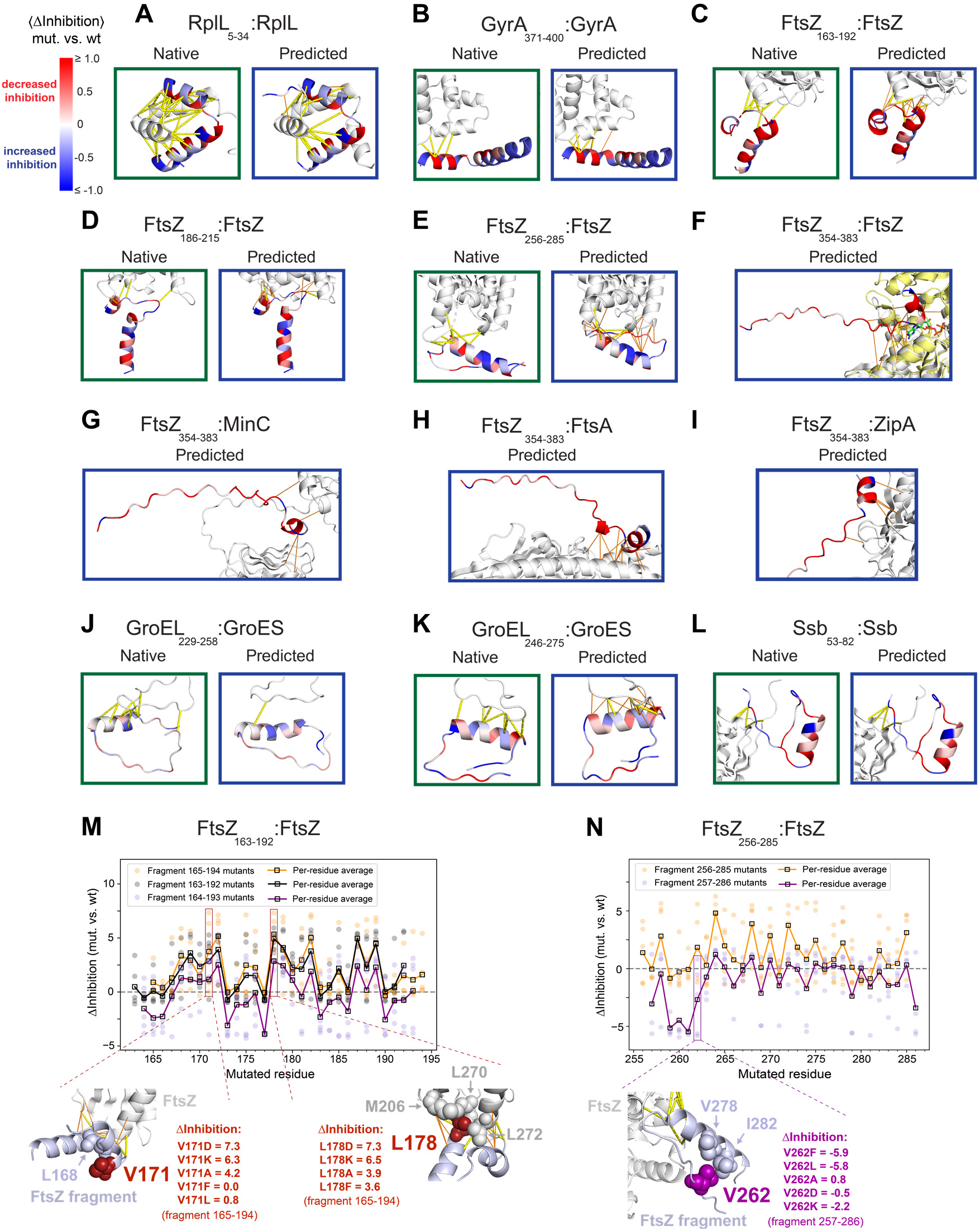
Deep mutational scanning of protein fragments reveals functional roles of interface-binding residues. (**A**) – (**L**) Fragment binding modes as per native full-length protein experimental structure (green boxes) and FragFold models of fragment binding (blue boxes) for each indicated inhibitory peak (FragFold model is of corresponding predicted peak fragment, as in Table 1), with the full-length protein target in white. Fragment residues are colored by residue-averaged ⟨Δ_Inhibition_⟩ from blue (mutations enhance inhibitory activity) to red (mutations weaken inhibitory activity) as indicated by the colorbar in (A). Native contacts are shown as yellow dashed lines and additional FragFold-predicted contacts are shown as thinner orange lines. For (E), the FragFold model of one FtsZ monomer is overlaid with the crystal structure of a FtsZ dimer from the filament structure (PDB ID 6unx) in yellow, and the structure of bound GTP in the active site is included. (**M**) – (**N**) *Above:* All mutations at each residue and their corresponding individual Δ_Inhibition_ values are plotted across several protein fragments derived from inhibitory peaks as indicated. *Below:* Example mutationally sensitive residue positions are selected to demonstrate structural model context of each position, and effects of mutating to each alternative residue included in the deep mutational scan at that position.

We found that residues directly involved in key predicted interactions commonly exhibit substantial sensitivity to mutation (**Fig. 6**, **Fig. S10**). Using positive average mutational sensitivity at each position as a metric (⟨Δ_Inhibition_⟩ > 0.5), we systematically assessed to what extent mutations at modeled folding and binding interaction residues disrupted inhibitory activity. Overall, across all inhibitory peaks, 54% of all FragFold-predicted protein-fragment contact residues (of 69 total) and 48% of FragFold modeled fragment folding contact residues (of 58 total) were detrimental to mutate (fragment ⟨Δ_Inhibition_⟩ > 0.5), compared to only 36% of residues overall. These values were statistically significant compared to a null model of random contact site distribution (*p* = 2.4 x 10^-3^ for the FragFold-predicted protein-fragment contacts, and *p* = 1.3 x 10^-3^ for predicted folding contacts; *Materials and Methods*). By way of comparison, structural models of fragment binding modes derived directly from native full-length protein structures were generally less aligned with the experimental data: 47% of native binding contacts and 41% of native folding contacts were detrimental to mutate, with corresponding *p*-values of 0.021 and 0.024, respectively. We found the statistical significance of the correlation between mutational sensitivity and predicted contact sites was not driven by bias in the mutational sensitivity of residue types (e.g., hydrophobic, charged, or polar; *Materials and Methods*), with *p*-values continuing to indicate high statistical significance when controlling for the residue type distribution of modeled contact sites in the null model (*p* = 1.5 x 10^-3^ for protein-fragment contacts predicted by FragFold, and *p* = 0.021 for those from native structure-based models). However, the modeled folding contacts were affected more substantially when controlling for amino acid type, with *p* = 0.059 for FragFold-predicted folding contacts and *p* = 0.35 for native structure-based models. Overall these results suggest that FragFold captures key aspects of how inhibitory protein fragments bind protein targets *in vivo*, with predicted models generally superior to models derived from the full-length protein interaction interface.

We further quantified the correlation of FragFold structural contact sites with positive mutational sensitivity (⟨Δ_Inhibition_⟩ > 0.5) for each inhibitory peak using additional metrics (**Table S3**). We found substantial true positive rates, true negative rates, and positive and negative predictive value statistics for most inhibitory fragment peaks. Overall, the Matthews correlation coefficient (MCC) between mutational sensitivity and FragFold-predicted contacts was positive (0.17) and highly statistically significant (*p* = 1.2 x 10^-4^ compared to the null model), indicating predictive value. On a per-fragment basis the MCC was generally positive (**Table S3**), with the exception of two fragment peaks: GroES fragment 226-255 and Ssb fragment 53-82 (MCC of ∼0, indicating no correlation). The deep mutational scanning data therefore provide support for most of the binding modes predicted, but not in the case of these two fragments. Strikingly, for the case of the GroES 226-255 inhibitory peak the MCC was much improved using the native structural model (**Table S3**, **Fig. 6*J***), suggesting that in this case the deviations from the native structure predicted by FragFold may be erroneous. (Indeed, this case stands out as one where FragFold predicts only one fragment-protein contact.)

Beyond the average effects of mutating each position of an inhibitory fragment, as described by ⟨Δ_Inhibition_⟩, the specific effects of each mutation at each residue, Δ_Inhibition_, varied widely. Individual values ranged from 7.3 to –6.2, a >2^13^-fold range in relative growth rate effects. Specific mutational effects on fragment inhibition could often be rationalized in terms of structural features of the fragments and their interactions. Here we consider two example inhibitory peaks exhibiting some of the strongest measured gain-of-function and loss-of-function mutational effects (**Fig. 6*M-N***).

In the case of residue V171 in FtsZ inhibitory fragment peak 163-192, the V171D mutation is highly detrimental to inhibitory function, presumably because it disrupts hydrophobic interactions with L168 that stabilize an alpha-helical structure (**Fig. 6*M****, left*). While mutations at this position to charged residues (D, K) or a small residue (A) are highly perturbative (e.g., Δ_Inhibition_ = 7.3, 6.3, 4.2 for D, K, and A in fragment 165-194), mutations to alternative hydrophobic residues (F, L) are well-tolerated (Δ_Inhibition_ = 0.0, 0.8). Similarly, FtsZ fragment residue L178 in the same inhibitory peak forms key interactions with full-length FtsZ in the FragFold structural model, positioned for hydrophobic interactions with L270, L272, and M206. Consistent with these hydrophobic interactions being important for fragment-protein binding, mutation of L178 to charged residues (D, K) is highly detrimental to binding (**Fig. 6*M****, right*). Fully consistent with our results, full-length FtsZ L178E (similar to L178D in mutant fragments) was previously found to strongly inhibit FtsZ filament assembly (48), with this variant only able to form filaments at crystallographic concentrations (32, 48).

Considering example mutations yielding *improved* inhibitory fragments, we find that variants V262F and V262L of FtsZ fragment 257-286 massively increase inhibitory activity (Δ_Inhibition_ = -5.9, -5.8; **Fig. 6*N***). V262 is near the base of a hairpin-like structure formed by the fragment, reminiscent of a helix-loop-helix motif, and is physically adjacent to hydrophobic residues V278 and I282 on the other side of the hairpin. Therefore, activating mutations V262F and V262L adding larger and more extended hydrophobic residues can be rationalized as providing stronger hydrophobic interactions with the other side of the hairpin base, stabilizing folding and therefore binding to the full-length FtsZ target (**Fig. 6*N***). Consistent with this interpretation, mutation to alanine does not provide this gain-of-function phenotype (Δ_Inhibition_ = 0.8), and nor does mutation to charged residue D (Δ_Inhibition_ = -0.5). Mutation to K intriguingly provides a partial benefit (Δ_Inhibition_ = -2.2), but more weakly compared to F or L at this position, perhaps due to interactions with the extended hydrophobic portion of the lysine sidechain.

Overall, we found that experimental high-throughput mutational scanning of protein fragments uncovers highly cooperative folding and binding of protein fragments; reveals that mutations to generate improved fragment inhibitors are readily accessible; and supports the binding modes predicted at scale with FragFold, further reinforcing that pervasive inhibitory protein fragments tend to act by inhibiting native protein interactions in cells (20).

## Discussion

In this work, we demonstrate the power of high-throughput AlphaFold predictions to discover protein interaction-inhibitory protein fragments and predict their binding modes. The results presented here are based on predictions and measurements for a panel of six diverse essential proteins for which binding partners are known. Yet the approach is expected to generalize to a much larger set of targets. Similar results are anticipated for any protein under conditions in which the relevant protein interactions become important for cell growth. In applying this approach to larger collections of proteins, knowledge of relevant interaction partners that could be the source of inhibitory fragments might come from genetic screens, pull-down experiments, colocalization measurements, or *in vitro* biochemistry, as tabulated in a number of databases (49–51).

An important parameter in evaluating our predicted binding peaks was the peak width – that is, the number of distinct overlapping protein fragments predicted to bind the target protein. In our binding prediction specificity tests, we observed narrower peaks for off-target compared to on-target interactions (**Fig. S4, S5**). We speculate that the exception to this trend, the fragment scan of RplL (**Fig. S5*C***), may reflect true binding promiscuity. Indeed, the apparently off-target interactions of RplL fragments with DNA gyrase subunits GyrA and GyrB may help to explain the outsized inhibitory effects observed for N-terminal fragments of RplL in cells (20). In general, binding peaks were more likely to match a measured inhibitory peak when the number of contributing protein fragments was sufficiently large, which we incorporated as a threshold in our automated pipeline for inhibitory fragment discovery (*Materials and Methods*). This filter based on predicted binding peak (cluster) width allowed us to remove false positive binding predictions, contributing to the ∼68% success rate in predicting inhibitory fragments systematically. This finding was reminiscent of other recent work showing that the number of overlapping protein sequence regions yielding comparable predictions improved confidence in protein-protein binding predictions using AlphaFold-Multimer (52), and more broadly the benefits of modeling short segments of protein sequence to predict binding interactions (52, 53).

It is instructive to compare the results from FragFold with other applications of AlphaFold to predict protein-peptide binding as well as full length protein-protein interactions. Previous work also using AlphaFold with monomer weights has demonstrated a ∼40–60% success rate in predicting known protein-peptide complex structures (24, 54); a similar ∼60% success rate was reported for correctly predicting pairwise protein-protein complexes (∼45% when using unpaired MSAs) (55). Our findings of ∼87% of known PPI-inhibitory fragments predicted to bind in a native-like manner and ∼68% of predicted binding peaks overall agreeing with measured inhibitory peaks compared well with these prior success rates. To the extent that our results reflect improved performance, we propose that this may arise partially from the benefits of deep MSAs available for protein fragments in contrast to peptides (24, 54), in the case of protein-peptide interactions. In addition, our results suggest that AlphaFold can be more successful at predicting interaction interfaces of full protein-protein complexes using short regions of sequence rather than the full-length sequences of both partners (e.g., **Fig. S6**). A similar conclusion was reached by Lee and colleagues, who found that AlphaFold-Multimer prediction accuracy of short linear motif (SLiM) binding to protein targets was substantially improved using only fragments of each binding partner, and decreased as the lengths of both protein partners was increased (52). Bret *et al.* and Yu *et al.* obtained similar results (53, 56). Our results suggest this trend should generalize outside of SLiM binding, and does not depend on training on protein complex structures as employed in AlphaFold-Multimer (as our predictions were made entirely using the monomer weights of the original AlphaFold2 model). In work that became available on *bioRxiv* while this paper was in revision, Mondal and colleagues also modeled peptides tiling across native proteins with an AlphaFold-based pipeline, across 12 protein-protein interactions, with the goal of predicting full-length protein-protein interaction binding epitopes (57). The authors found that previously determined native epitopes were the top-ranked binders in 7/12 instances, and lower-ranked (3/12) or undetected (2/12) in the remainder of cases. Overall, the authors’ observed rate of 10/12 (∼83%) native epitope peptides detected as significant binders is congruent with our finding that, using FragFold, 87% of known interface inhibitory peptides are successfully predicted.

We also found that the number of fragment-target contacts from AlphaFold structural models (*N*_contacts_) weighted by the predicted interface TM score (ipTM) (30, 58) was a strong predictor of inhibitory protein fragments. This approach, placing the structural model first and the prediction confidence of AlphaFold second, stands in contrast to other approaches that evaluated predictions purely based on ipTM or pLDDT (24, 52, 59). The demonstrated effectiveness of weighted *N*_contacts_ in the context of inhibitory protein fragment prediction suggests it may be a useful metric for other applications as well. Our finding of the predictive value of *N*_contacts_ is in agreement with the recent work of Bryant and colleagues. Similar to our weighting approach, they found that a combination of pLDDT and *N*_contacts_ was a potent discriminator (55).

The automated overlap analysis of predicted binding clusters (revealing predicted binding peaks) and measured inhibitory peaks had some limitations. First, the floor for successful prediction was high due to the biology of inhibitory protein fragments of this length: the *f* _predicted binding clusters inhibitory_ expected from random placement of peaks was 34 – 66% (51% across all proteins) due to the significant fraction of protein sequence corresponding to inhibitory peaks (20). Second, the maximum success rate is limited by several factors: not all predicted binding modes from AlphaFold represent true positives, not all binding modes are inhibitory, and not all possible cellular protein binding partners of each protein fragment are considered in the predictions. These challenges are exemplified by the predicted binding peak centered around FtsZ residue 60 (**Fig. 2*A***, *left*, blue), which has no corresponding inhibitory activity (**Fig. 2*A***, *left*, green). In this light, the obtained values of *f* _predicted binding clusters inhibitory_ (68%) and *f* _inhibitory peaks predicted_ (64%) represented successful, but imperfect, predictions. Another limitation is that although predicted binding clusters overlapped with measured inhibitory peaks, the predictions and measurements often exhibited an offset (**Fig. S2**). This indicates that several nearby tiling fragments should be tested for each FragFold-predicted binding peak to identify functional inhibitors. Reducing this offset between the predictions and the experiments represents an area for future improvements and may reflect limitations in the precision of binding predictions.

In interpreting deep mutational scanning data on protein fragments as obtained in this work, we caution that noncontact residues may also influence fragment folding and therefore protein binding, contributing to the mutational sensitivity seen in noncontact sites (**Fig. 6**). Such effects are likely particularly prevalent under cooperative folding and binding. We also note that our data do not distinguish between potential mutational effects on protein fragment abundance (which in turn influences binding) and effects directly on binding, which may introduce false negatives for loss-of-function effects in particular. Finally, the mutational sensitivities measured likely reflect additional binding interactions not fully captured by the models. Nonetheless, the correlations seen between our deep mutational scanning results and FragFold predictions strongly suggest that FragFold is capturing key aspects of the binding interactions of these inhibitory peptides *in vivo*.

Our results suggest multiple applications for FragFold. First, the high success rate of our predictions (∼68% of predicted binding clusters corresponding to experimentally measured inhibitory peaks) shows that using FragFold as a computational screen for fragment-based inhibition should be fruitful across diverse proteins of interest. Such predictions have direct applications and are highly synergistic with high-throughput experimental measurements of fragment-based inhibition (17–20). The revolution in DNA synthesis capabilities (60, 61), combined with the ongoing revolution in high-throughput sequencing, has enabled these high-throughput experimental approaches (19); nonetheless, limitations persist in library size and cost. In applications with the primary goal of discovering novel inhibitory peptides, e.g., as leads for novel therapeutics, or probes for systematically disrupting PPIs, pre-screening for likely inhibitory protein fragments with FragFold stands to substantially reduce the protein sequence space that needs to be scanned. This will allow for the coverage of significantly larger sets of fragments in a library of given size, greatly accelerating inhibitory fragment discovery and large-scale systematic characterization. FragFold should likewise enable the economical use of smaller libraries for inhibitor characterization. In future work, FragFold has the promise to further synergize with additional experimental modalities, including cell-display approaches and massively-parallel assays of folding and binding interactions (62–66). Indeed, we note that FragFold specificity was generally good for the protein fragments tested (on the order of ∼10-fold) but that the raw number of off-target predictions will nonetheless likely grow faster than the number of on-target predictions when the proteome scale is considered. Thus, experimental information on likely interaction partners will be a powerful partner to FragFold in predicting fragment-protein interactions at scale.

Second, FragFold can complement high-throughput measurements by supporting the interpretation of peptide inhibitory mechanisms at scale. We demonstrate this complementarity in this work by combining FragFold structural modeling with high-throughput measurements of inhibitory function *in vivo*, both providing further support for the interpretation of the measured inhibitory effects as driven by dominant-negative competitive inhibition of native interactions (ref. (20); **Table 1**), and predicting novel FtsZ IDR fragment binding modes in agreement with orthogonal functional data. We also demonstrate the synergy of experimental deep mutational scanning of protein fragments and prediction of fragment binding modes, including for the interpretation of variants that improve inhibitory activity, illuminating a promising route towards peptide drug development. Our findings suggest that AlphaFold has learned generalizable principles of protein fragment binding that can be applied to discover novel interactions. In this, our results add to a growing body of work showing successful AlphaFold prediction of previously unknown protein-protein interactions aligning with experimental measurements. Examples include a phage defense protein complex structure concordant with hydrogen-deuterium exchange mass spectrometry measurements (67), and a Kelch domain pocket – SLiM interaction in agreement with BRET measurements and mutagenesis results (52). Other recent work has similarly leveraged the synergy of AlphaFold predictions with high-throughput experiments, including the combination of mass spectrometry-based protein interactomics with AlphaFold modeling (59).

In conclusion, we have demonstrated the power of AlphaFold to systematically predict protein fragments that inhibit native protein interactions in living cells. The approach we have developed to scale these predictions across tiling protein fragments, FragFold, promises to be broadly applicable to the discovery of novel inhibitory peptides across proteomes. This is supported by our ability to predict nativelike inhibitory fragments across diverse proteins, as well as previously unknown binding modes of an intrinsically disordered region. FragFold enables both screening for and systematic characterization of inhibitory fragments and provides a natural complement to high-throughput experimental methods for measuring peptide function.

## Materials and Methods

### Selection of proteins to computationally scan with fragments

Proteins computationally fragmented and investigated with FragFold were selected from among those experimentally measured with a high-throughput approach in previous work (20). We chose to investigate fragments of FtsZ, RplL, GyrA, Ssb, LptG, and GroEL because each forms structurally characterized protein-protein interactions that we might expect AlphaFold to be able to predict (as opposed to interactions of unknown structure, protein-nucleic acid interactions, or interactions with folding intermediates).

### Predicting fragment-protein complex structures

We used ColabFold (version 1.5.2) to predict the structure of protein fragments interacting with full-length proteins. We used the AlphaFold2 monomer weights (`model_type = monomer_ptm`) with proteins and fragments modeled as separate chains, as natively implemented in ColabFold via the relative positional encoding (29), to ensure that no information on protein-protein interactions was utilized. No templates were used. Other settings were set to ColabFold defaults. We modified the protocol for multiple-sequence alignment (MSA) generation to maximize the depth of the MSA for the fragments. Specifically, we used MMseqs2 (68) to construct MSAs individually for all full-length proteins from which fragments were derived. The residues in the MSA corresponding to the fragment were then extracted from the full-length MSA and used to construct a fragment MSA (for example, for a 30-amino acid fragment starting at residue 100, we extract all amino acids from sequences aligned to positions 100-129 in the full-length MSA). We then concatenated the fragment and protein MSAs, as required by ColabFold as the standard input for a multi-chain prediction. Specifically, we performed a row-wise concatenation of the full-length and fragment MSAs. This yielded unpaired MSAs, where each row contains a single sequence, mapping to either the full-length protein or the fragment, and gap tokens for the other chain. We used unpaired MSAs to avoid co-evolutionary signals of protein-protein interactions being used in the prediction. Note that while the MSAs are concatenated (as required by ColabFold), the sequences of the full-length and fragment chains are distinct and not connected. We iterated this process for all fragments of full-length proteins to generate custom input MSAs for ColabFold. The runtime was highly dependent on the total number of residues in the fragment and full-length protein chain. For example, using 4 A100 GPUs it took 16 hours to predict the structures of 354 FtsZ fragments (30 aa) with full-length FtsZ (306 residues) whereas it took 210 hours to predict the structures of 843 GyrA fragments (30 aa) with full-length GyrA (872 residues).

### Native interface recovery

We compared experimentally determined structures of interacting full-length proteins (i.e., native) with ColabFold-predicted structures of fragments and full-length proteins (i.e., predicted). These comparisons were performed for a set of known inhibitory protein fragment peaks from previous experimental measurements that mapped to structurally characterized protein-protein interaction interfaces; these peaks were identified previously (20) or in this work based on inspection of the inhibitory activity profile of high-throughput fragment scans and corresponding experimental structures of protein complexes. For each inhibitory peak, we selected a representative protein fragment possessing the maximal inhibitory activity of all fragments in the peak. The experimental structure for this representative inhibitory fragment – protein interaction was compared to a corresponding AlphaFold-predicted model for a fragment of similar or identical sequence position (offset of 0 – 13 residues, see **Table 1**) corresponding to a local maximum of weighted *N*_contacts_. The top AlphaFold model ranked by weighted *N*_contacts_ was used. After defining all residues corresponding to the fragment of interest in the experimental structure, we removed all other residues from the chain to create a model of the fragment interacting with the full-length protein chain as it does in the context of the full protein-protein interaction. We compared the native and predicted structures using contact recovery and backbone RMSD metrics.

We defined interface residue-residue contacts between the fragment and full-length protein chain based on heavy-atom distances. Specifically, a contact was defined between two residues if they had at least one pair of heavy atoms within 4.0 Å. We defined interface contacts in the native structure and calculated the native contact recovery as the fraction of native contacts observed in the predicted structure.

Peptide root-mean-square deviation (RMSD) was calculated between backbone atoms (N, Ca, C, O) of the native and predicted structures, after optimal superposition of the full-length protein chain using the Bio.PDB Python package. In some cases, residues were missing from the native structure and the number of residues used in the calculation was less than the number of residues in the fragments.

Interface RMSD was calculated between all interface residues in the native structure and corresponding residues in the predicted structure. We designated interface residues using the earlier defined contact definition, with a larger distance cutoff of 8.0 Å. The optimal superposition between the native interface residues and corresponding residues in the predicted structure was applied and the RMSD was calculated. If there were fewer than 5 corresponding residues in the fragment or full-length protein chain, we did not compute the RMSD.

### Automated prediction of inhibitory peaks

We filtered and clustered all of the predicted complex structures to predict potential inhibitory peaks. For each set of predicted structures, i.e., ColabFold predictions of fragments of one protein interacting with a full-length protein, we filtered predictions by multiple criteria. Predicted structures were only considered if they had at least 3 interface contacts within 3.5 Å, 3 ipTM-weighted interface contacts, and an ipTM value of 0.3 or greater. The predicted structures (up to 5 per fragment) were then clustered using Tanimoto distance (69, 70) over interface contacts. Specifically, we performed hierarchical complete-linkage clustering with a maximum Tanimoto distance of 0.6. We then filtered the clusters based on the number of unique fragments in the cluster, requiring at least 6 fragments per cluster, and merged heavily overlapping clusters to obtain the final set of predicted binding clusters. We used the set of all residues in a cluster (from any fragment) to constitute a predicted inhibitory peak. We obtained the final set of parameters by scanning over potential values for minimum *N*_contacts_, weighted *N*_contacts_, ipTM, and number of overlapping fragments per cluster, as well as maximum Tanimoto distance. We manually inspected the predicted clusters and optimized for the resulting *f* _inhibitory peaks predicted_, *f* _predicted binding clusters inhibitory_, and a number of predicted peaks called similar to the number of experimental peaks measured. The minimum cluster size of 6 was selected based on this optimization of *f* _inhibitory peaks predicted_ and *f* _predicted binding clusters inhibitory_ (**Fig. S3)**. Matching of predicted binding peaks to inhibitory activity peaks (called from the experimental data as described below) was assessed based on peak overlap, with an overlap of at least 2/3 of the fragment length (i.e., 20 aa for 30 aa fragments) required to consider two peaks (widths ranging from 30 − 67 aa for 30 aa fragments) to match.

### Calling inhibitory peaks from experimental data

We automatically annotated peaks in the experimental inhibitory data using a simple protocol. Measured *in vivo* inhibitory effects were expressed as the enrichment (*E*) of protein fragments in a high-throughput massively parallel growth selection experiment, such that negative *E* values corresponded to inhibitory activity (20). We filtered out fragments that had inhibitory effects less than a gene-specific inhibitory effect cutoff corresponding to an enrichment Z-score of −2.5. The only exception to this universal Z-score cutoff was made for the protein GroEL, in which case we used a cutoff corresponding to a Z-score of −2 due to the apparent lower signal:noise ratio of the inhibitory peaks previously identified (20). After filtering by this Z-score cutoff, only inhibitory fragments remained. To group these fragments into peaks, we considered each fragment in turn, proceeding from the N- to the C-terminus. The N-terminal most inhibitory fragment was considered the starting point of the first peak; for each subsequent inhibitory fragment, the fragment was added to an existing peak if the N-terminal residue of the new fragment was less than 5 aa from the N-terminal residue of the nearest inhibitory fragment in the peak and no more than 25 residues away from the N-terminal residue of the first fragment in the peak. Otherwise, a fragment was considered the starting point for a new peak. This simple algorithm successfully reproduced all inhibitory peaks called by hand here and in previous work (**Table 1** and ref. (20)); comparing manual and automatically-called peaks, we note that in a few cases two nearby peaks were merged, and in other cases peaks were split into nearby sub-peaks.

### Peak prediction null model

We developed a null model for simulating peak prediction with randomized peak positions to obtain null expectation values and *p*-values for the sensitivity (*f* _inhibitory peaks predicted_) and specificity (*f* _predicted binding clusters inhibitory_) statistics that we measured for our FragFold results. For each protein, we assumed the same number of predicted peaks as calculated by FragFold, and assigned peak positions at random, with starting residues sampled uniformly from the range [1, *L* – *k*) (where *L* and *k* are the amino acid lengths of the protein and fragments, respectively). Randomly placed peaks that overlapped another randomly placed peak with ≥70% sequence overlap were rejected and resampled. We set the width of each null model peak to the mean peak width from FragFold across all predictions over all proteins. *f* _null, inhibitory peaks predicted_ and *f* _null, predicted binding clusters inhibitory_ were then calculated for each protein and across all proteins based on overlap with the fixed inhibitory peaks from experimental data (20) just as for the FragFold results. We repeated this 5,000 times for each protein to obtain distributions that we used to calculate null expectation values (means of *f* _null, inhibitory peaks predicted_ and *f* _null, predicted binding clusters inhibitory_ across all simulation runs), and *p*-values across all proteins for the FragFold results compared to this null model.

### Rosetta relaxation analysis of protein fragment and peptide binding modes

We used Rosetta Relax to perform local all-atom refinement (71, 72) of the structures predicted by FragFold as well as experimental structures of peptides bound to protein targets from the PepPro database (45). We constrained relaxation to the starting coordinates with the argument “-constrain_relax_to_start_coords". We controlled the strength of the constraint by modifying the relax script “InterfaceRelax2019”. We used the default script as well as a more constrained version where the value of coord_cst_weight was fixed to 1.0. We used the Rosetta interfaceAnalyzer with default settings to compute the dG_separated/dSASAx100 score (73). In all applications of Rosetta in this work, we used the standard weights, REF15, and Rosetta 3.13 Linux Release (2021.16.61629).

### Peptiderive predictions of protein fragment binders

Peptiderive predictions of protein fragment binders derived from experimental structures of protein-protein interaction interfaces were performed using the ROSIE Peptiderive server (46, 74). Predictions were run using PDB structures 6unx for FtsZ, 1rqu for RplL, 6rks for GyrA, 1eqq for Ssb, 1aon for GroEL, and 6mi7 for LptG, and default Peptiderive parameters. Peptide fragment interface scores and sequence positions were extracted from the Peptiderive output and compared to experimentally measured inhibitory effects and FragFold predicted binding across full protein sequences.

### Deep mutational scanning of protein fragments

We generated DNA templates encoding protein fragment sequence variants with 5 possible alternative residues included at each position in each fragment tested: alanine, leucine, phenylalanine, aspartate, and lysine. Alternative residues for each position thus included a small or bulky hydrophobic, an aromatic residue, and the insertion of a positive charge or a negative charge. For a subset of fragments, we also included 2 additional alternative residues, serine and proline. This led to at least 4, and up to 7, possible mutations at each position of each fragment, depending on the wild-type residue. Wild-type and mutant protein fragment coding sequences were then cloned into the pET-9a expression vector (Novagen) in an identical manner to previous libraries of protein fragments (20).

The fragment-encoding plasmid library was then transformed into electrocompetent *E. coli* BL21 (DE3) cells (Sigma-Aldrich) at >170-fold coverage of the library size. After 1 hour of recovery post-transformation with the library, cells were back-diluted into LB media (Gibco) supplemented with kanamycin (selecting for the pET-9a backbone) and 10 uM IPTG (inducer for fragment expression). Cells were then grown to an OD_600nm_ of 0.7, back-diluted into fresh LB supplemented with kanamycin and IPTG, and grown further overnight. Cells were then harvested, and plasmids were extracted from each sample (as well as the plasmid library input) via minipreps (Qiagen). Amplicons for high-throughput sequencing were generated in the same manner as previously (20), and paired-end sequencing was performed on an Illumina Nextseq platform. These measurements allowed the determination of fragment frequencies (*f*) in the population before and after selection and hence the inhibitory activity (enrichment, *E*) = log_2_(*f*_initial_ / *f*_final_). These experiments were performed in triplicate (3 biological replicates) and results from this experimental assay were highly reproducible as reported previously (20); the reported values of *E* for each fragment are the average across all experiments.

For each mutation at each position of mutagenized protein fragments, the mutational sensitivity Δ_Inhibition_ was determined as Δ_Inhibition_ = *E*_mutant_ – *E*_wt_, where –*E* is the inhibitory activity (enrichment). The threshold value ⟨Δ_Inhibition_⟩ > 0.5 was selected for systematic analyses of mutational effects as it visually selected the observed peaks (**Fig. S10**) and optimized both the enrichment for mutational sensitivity at contact sites, compared to all residues, and the average Matthews Correlation Coefficient (MCC) values for the correlation between inhibitory effects and mutational sensitivity. In order to determine *p*-values for the significance of the fractions of modeled contact sites that are mutationally sensitive, a null model of random site distribution was constructed by randomly shuffling the alignment between experimentally measured mutational sensitivity values and model binding or folding contact residues within each protein fragment 50,000 times. *p*-values were then calculated from this null model distribution as the fraction of shuffled results yielding a mutationally sensitive fraction at least as high as the experimentally determined value. Additional *p*-value calculations with control for the residue type were performed in the same way as described above with 10,000 shuffles of the alignment of contact residues and mutational sensitivity values, but this time requiring each shuffle to maintain the same frequency of residue types making contacts in the original unshuffled model, with types defined as follows: hydrophobic (A, F, L, I, M, V, W, Y); negatively charged (D, E); positively charged (K, R, H); polar (N, Q, S, T); and other (G, C, P).

## Acknowledgements

We thank S. Fields, C. Sanchez, R. M. Garner, P. L. Freddolino, H. K. Wayment-Steele, S. Ovchinnikov, and members of the Li lab for helpful discussions. A.S. was supported by NRSA fellowship F32 GM134557 from the National Institutes of Health. This work was supported by awards K99 GM148718 and F32 GM134557 (to A.S.), R35 GM124732 (to. G.W.L.) and R35 GM149227 (to A.K.) from the National Institutes of Health. We acknowledge the MIT SuperCloud for providing computational resources (75), and the MIT BioMicro Center for DNA sequencing.

## Author contributions

Research conceptualization and design were performed by A.S. and S.S. Design and implementation of the FragFold computational method was performed by S.S. with feedback from A.S. Fragment deep mutational scanning experiments were designed and performed by A.S. Data analysis was performed by A.S. and S.S., with suggestions from A.K. and G.W.L. The manuscript was written by A.S. and S.S. with feedback and revisions from A.K. and G.W.L.

## Data and code availability

The code used in this paper – and to run FragFold on any protein of interest – is available on GitHub (https://github.com/swanss/FragFold). Data are provided in the main text, the Supplementary Information, and a Source Data file available via figshare (doi: 10.6084/m9.figshare.24841269; URL: https://figshare.com/articles/dataset/Source_Data_for_Savinov_and_Swanson_et_al_2023/24841269).

## Supplementary Information

### Supplementary Figures

**Figure S1.**
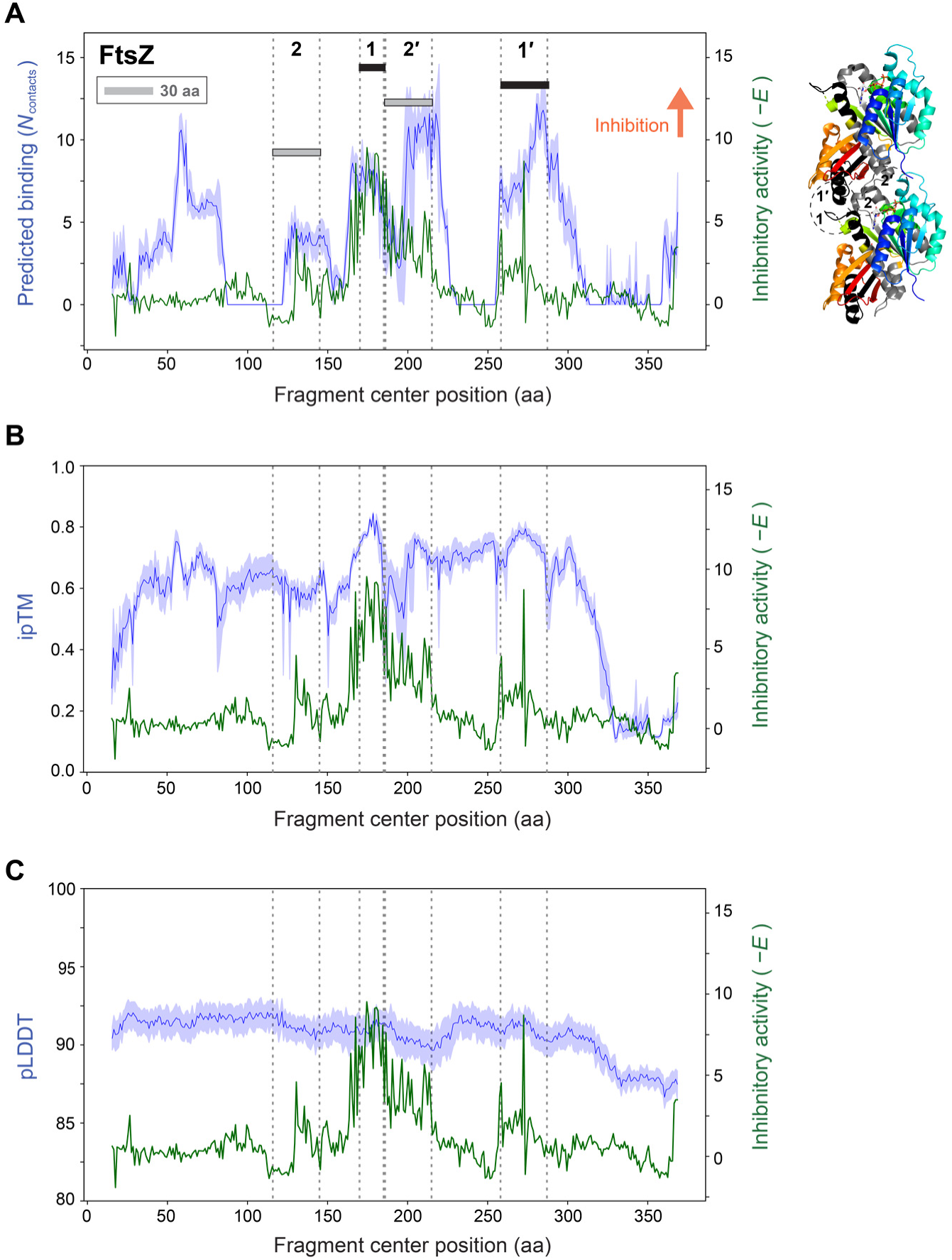
Unweighted *N*_contacts_ predicts inhibitory fragment peaks; AlphaFold confidence metrics alone do not. (**A**) As in Figure 2*A*, but with *N*_contacts_ (unweighted) instead of ipTM-weighted *N*_contacts_ shown. *Left*: Predicted target protein binding from FragFold, unweighted *N*_contacts_ (blue curves) vs. experimentally measured *in vivo* inhibitory activity (green curves, from ref. 20) for 30 aa fragment scans across *E. coli* FtsZ. Darker line and lighter outline for predicted binding data indicate mean and 95% C.I., respectively, across the 5 top-ranked structural models generated by AlphaFold. Filament formation interface regions **2**, **1**, **1′**, and **2′** are indicated. *Right*: crystallographic structure of two adjacent monomers of the FtsZ filament (PDB ID 6unx, ref. 32). Protein is colored from blue to red from the N- to the C-terminus. Regions corresponding to interface regions are annotated and highlighted in the same color scheme as in the fragment scan plot (*Left*). (**B**) As in (A), but ipTM is shown in blue. (**C**) As in (A), but pLDDT is shown in blue.

**Figure S2.**
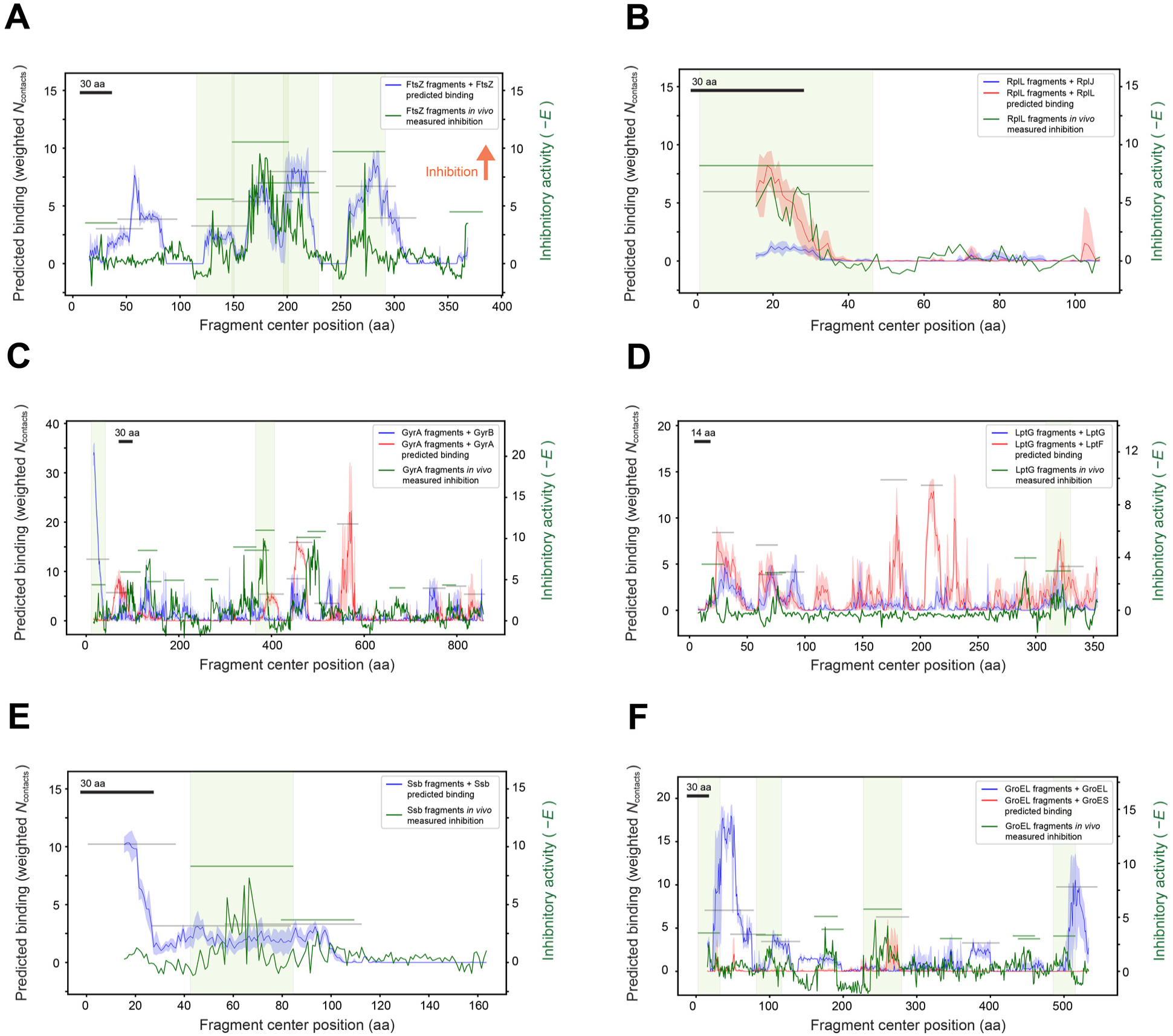
Automated peak discovery systematically reveals matching peaks between experimental inhibitory effects and predicted binding data. Predicted target protein binding (red, blue curves) from FragFold vs. experimentally measured *in vivo* inhibitory activity (green curves, from ref. 20) for fragment scans across 6 diverse *E. coli* proteins. Green highlights indicate peaks mapping to previously identified protein-protein interactions with corresponding inhibitory protein fragments (Table 1 and ref. 20). Darker line and lighter outline for predicted binding data indicate mean and 95% C.I. across the 5 top-ranked structural models from AlphaFold. Green and grey bars indicate peaks called by the peak-calling algorithm: green bars, inhibitory peaks from experimental data; grey, predicted binding clusters from FragFold results. Fragment scan results are shown for (**A**) FtsZ, (**B**) RplL (L7/L12), (**C**) GyrA, (**D**) LptG, (**E**) Ssb, and (**F**) GroEL. 30 aa fragments were used for all proteins except LptG, which was scanned with 14 aa fragments. aa, amino acids. Fragment sizes are also indicated by the black scale bar in each panel.

**Figure S3.**
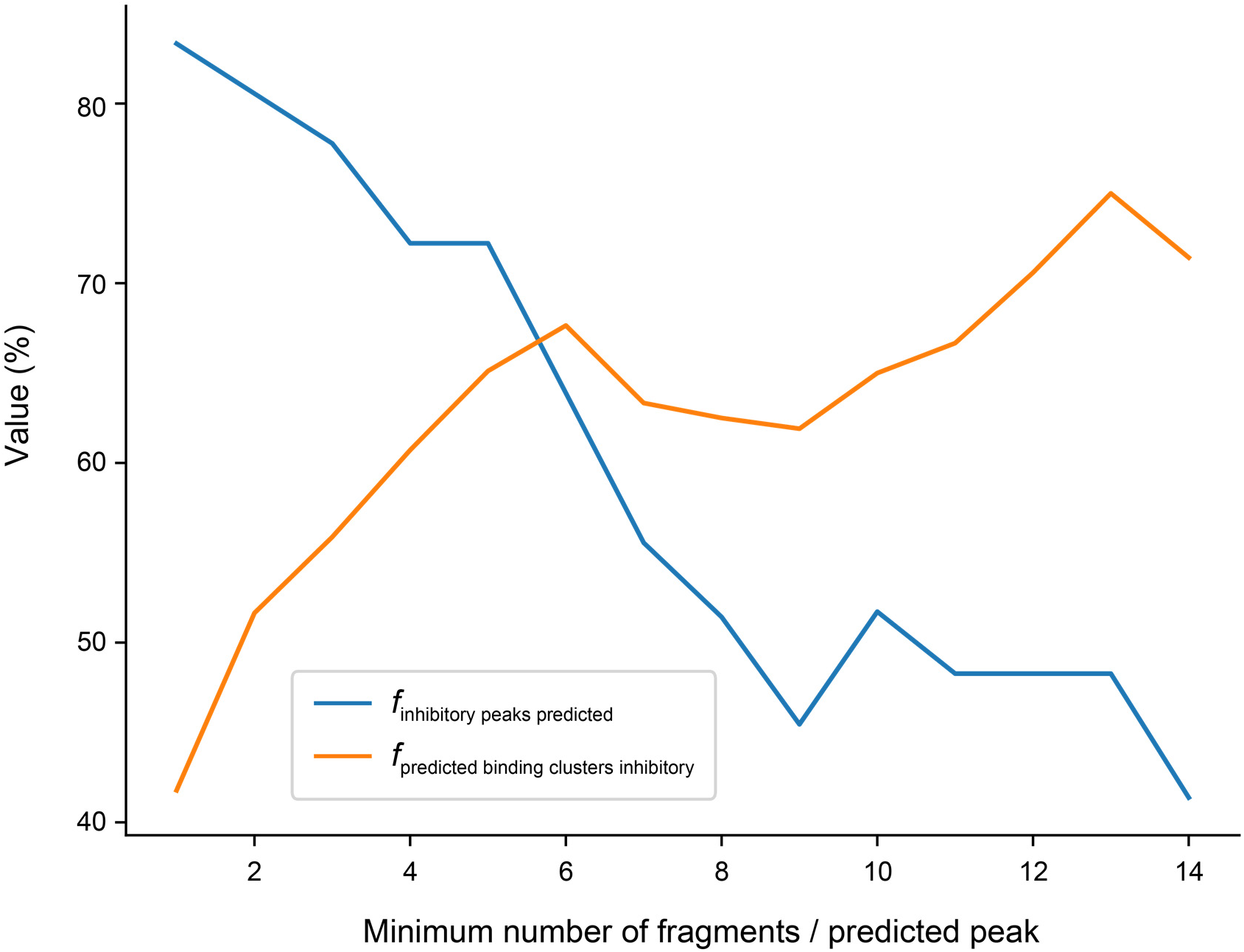
The minimum predicted peak width for automated inhibitory peak prediction was selected based on an optimum of sensitivity and specificity. Sensitivity (*f* _inhibitory peaks predicted_) vs. specificity (*f* _predicted binding clusters inhibitory_) plot is shown as parametrized by the minimum number of fragments per predicted binding peak (Tanimoto cluster size) accepted by the peak-calling algorithm.

**Figure S4.**
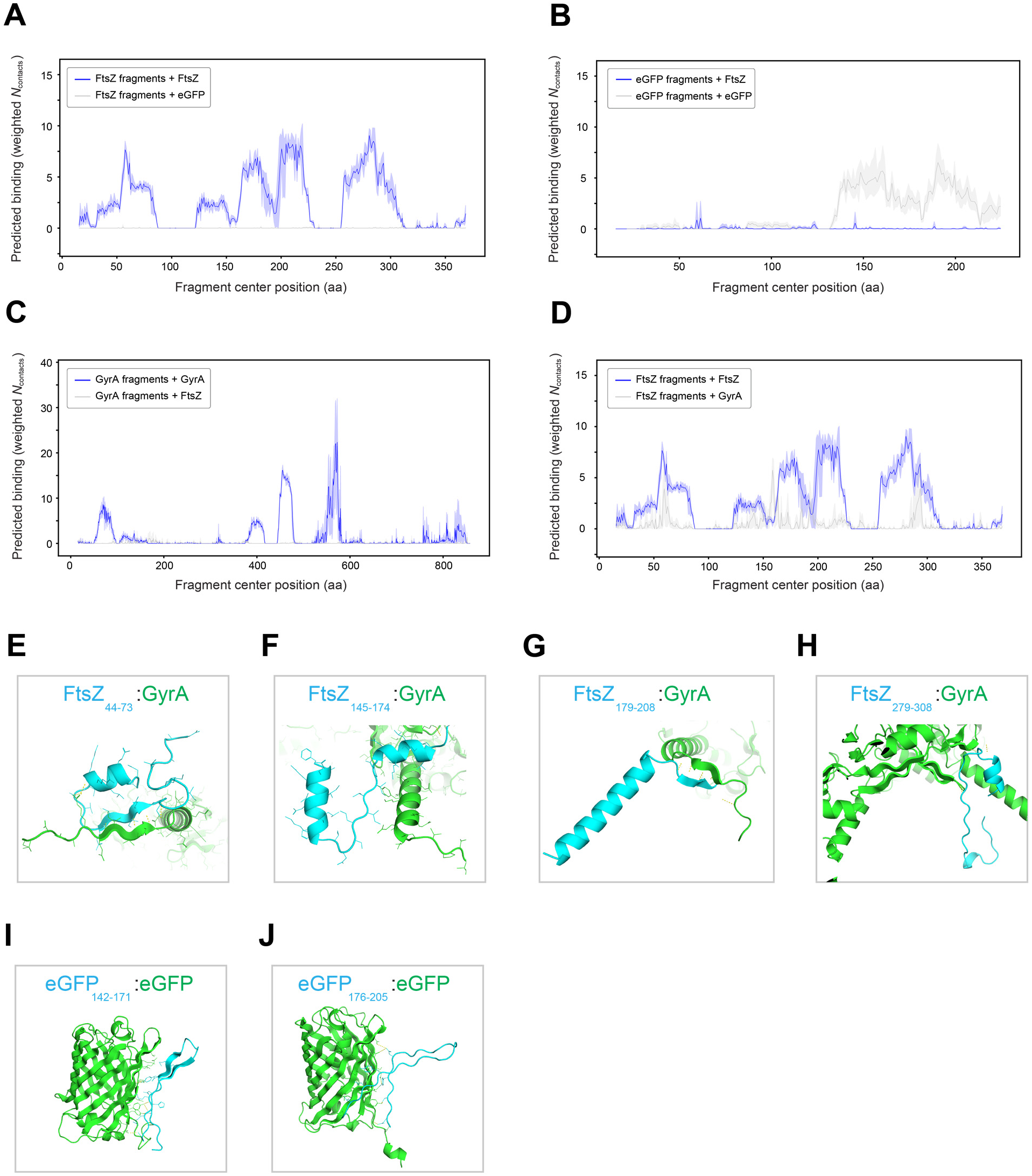
AlphaFold predictions of fragment-protein binding exhibit specificity. **(A)** – **(D)** Predicted target protein binding from FragFold for 30 aa fragment scans + target protein, depicted as in Figure 2. Colors are as indicated: (**A**) Blue, fragments of FtsZ + FtsZ; grey, fragments of FtsZ + eGFP; (**B**) Blue, fragments of eGFP + FtsZ; grey, fragments of eGFP + eGFP; (**C**) Blue, fragments of GyrA + GyrA; grey, fragments of GyrA + FtsZ; (**D**) Blue, fragments of FtsZ + FtsZ; grey, fragments of FtsZ + GyrA. aa, amino acids. (**E**) – (**H**) AlphaFold structural models of indicated fragments of FtsZ (cyan) + GyrA (green), corresponding to grey peak regions from (D). (**I**) – (**J**) As (E) – (H), but for fragments of eGFP + eGFP, corresponding to grey peak regions from (B).

**Figure S5.**
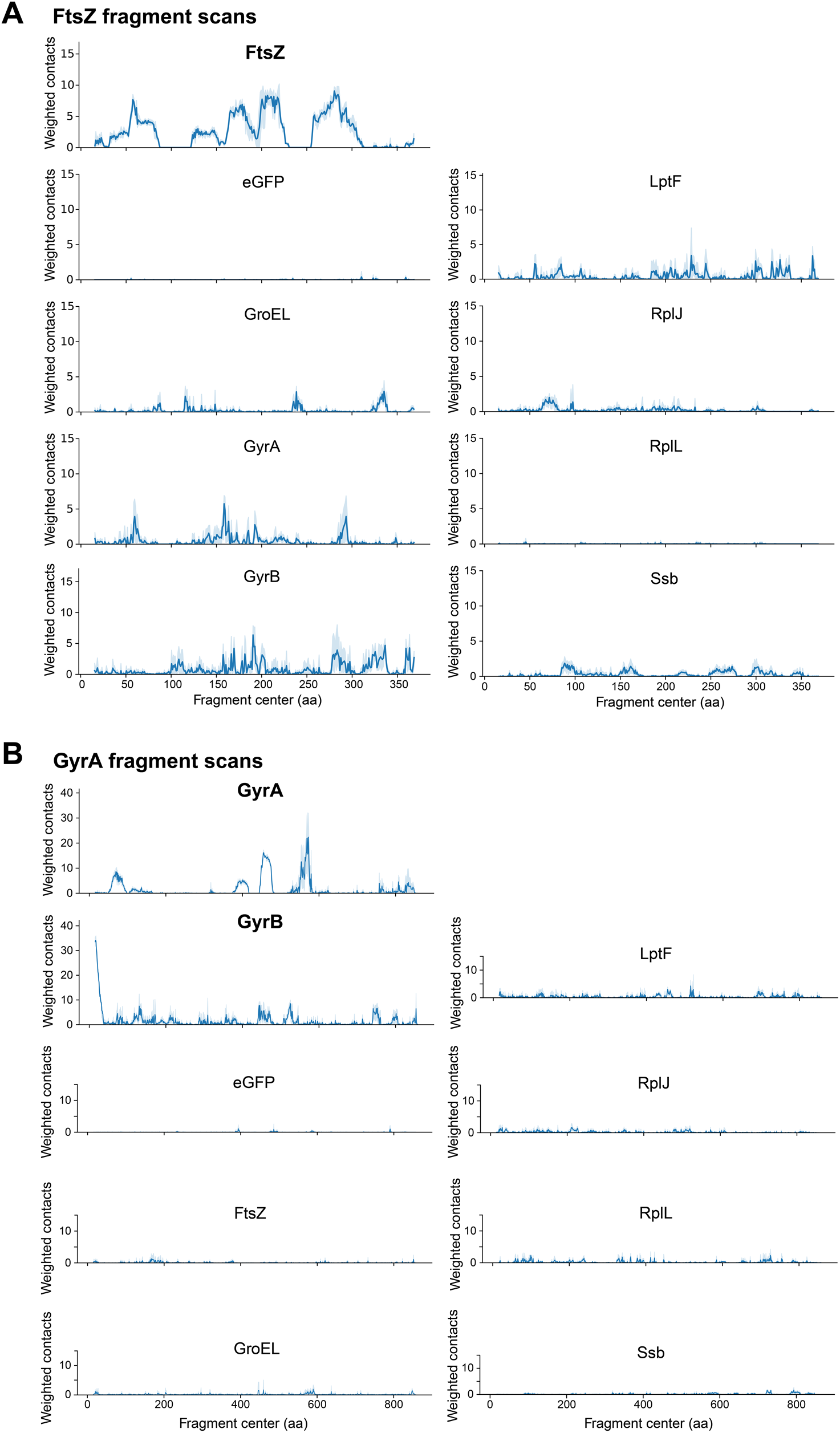
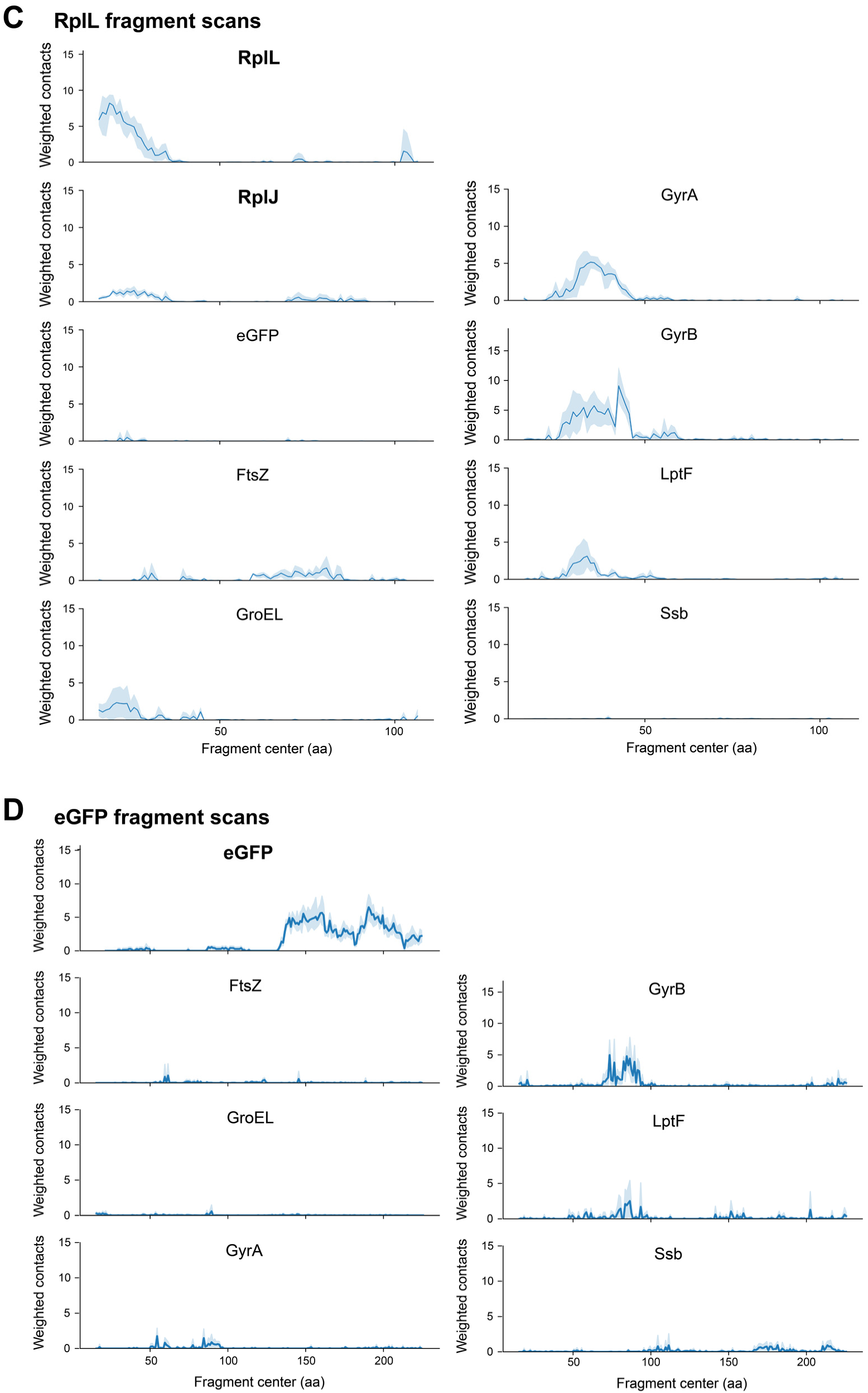
Further investigations of FragFold specificity. **(A)** – **(B)** Predicted target protein binding from FragFold for 30 aa fragment scans + target protein, depicted as in Figure 2, for on- and off-target binding partners, for fragments scanning across FtsZ (**A**) and GyrA (**B**) **(C)** – **(D)** As (A) – (B), but for fragments scanning across RplL (**C**) and eGFP (**D**)

**Figure S6.**
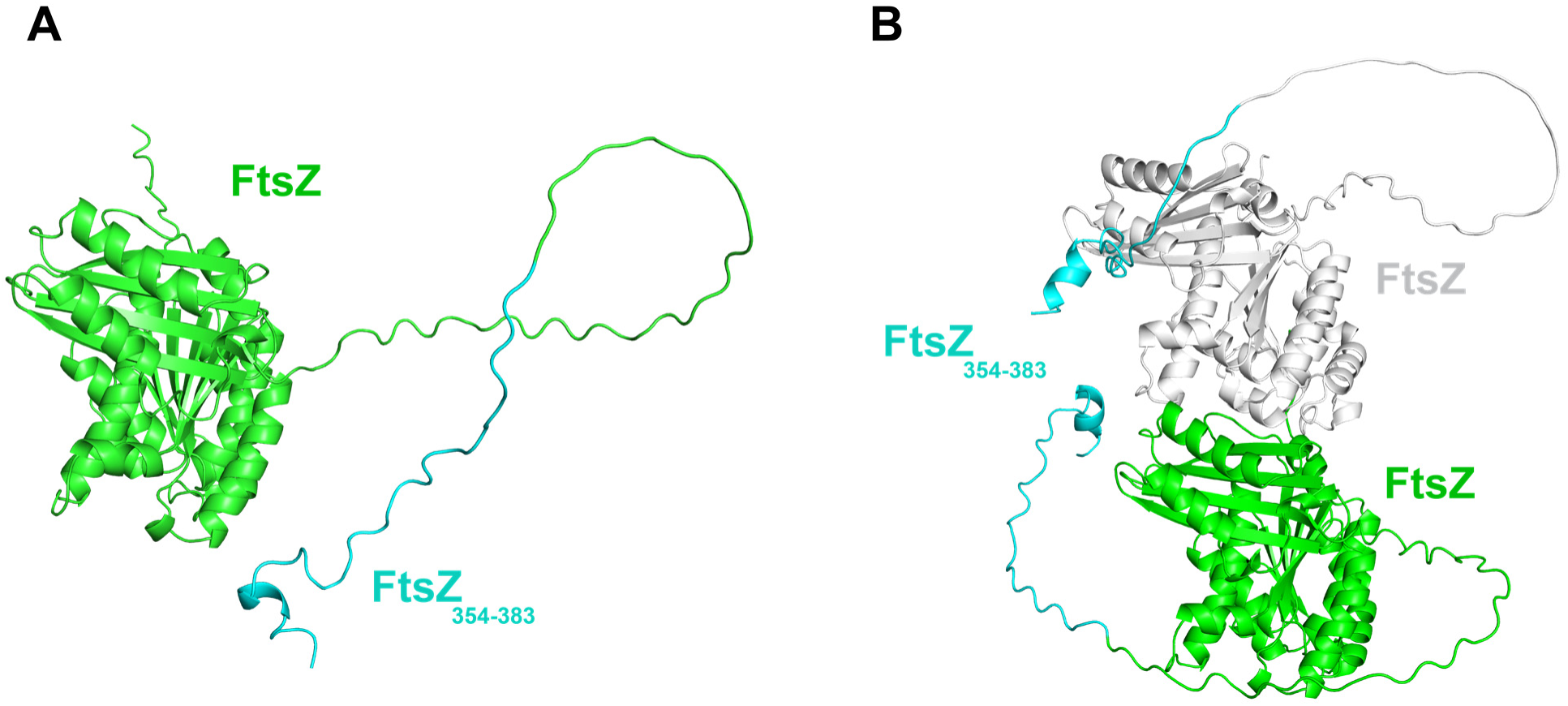
AlphaFold predictions for full-length FtsZ folding and dimerization fail to predict the C-terminal tail interaction revealed by FragFold. AlphaFold-predicted structures for full-length FtsZ (green and white), either as a monomer (**A**) or a dimer (**B**), with the C-terminal tail fragment region found by FragFold to bind at the GTPase active site (see Figure 4*B*) highlighted in cyan.

**Figure S7.**
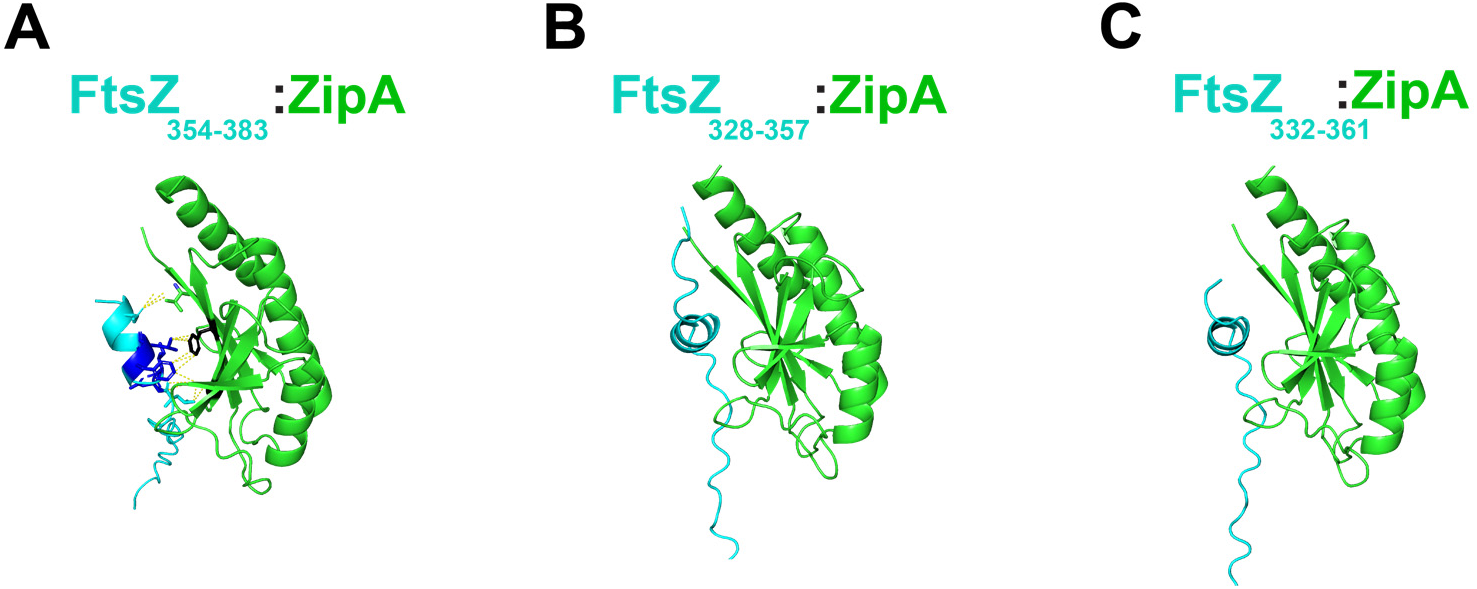
Additional FtsZ C-terminal tail fragments are predicted to bind the same face of ZipA. AlphaFold-predicted structures for indicated fragments of FtsZ (cyan) + ZipA (green). (**A**) Previously identified interacting regions on FtsZ (blue) and ZipA (black) are shown, as in Figure 4*E*, with residue-residue contacts shown as dashed yellow lines. (**B**) – (**C**) AlphaFold models for FtsZ fragments corresponding to FtsZ IDR peaks outside of the extreme C-terminal region (see Figure 4*A*).

**Figure S8.**
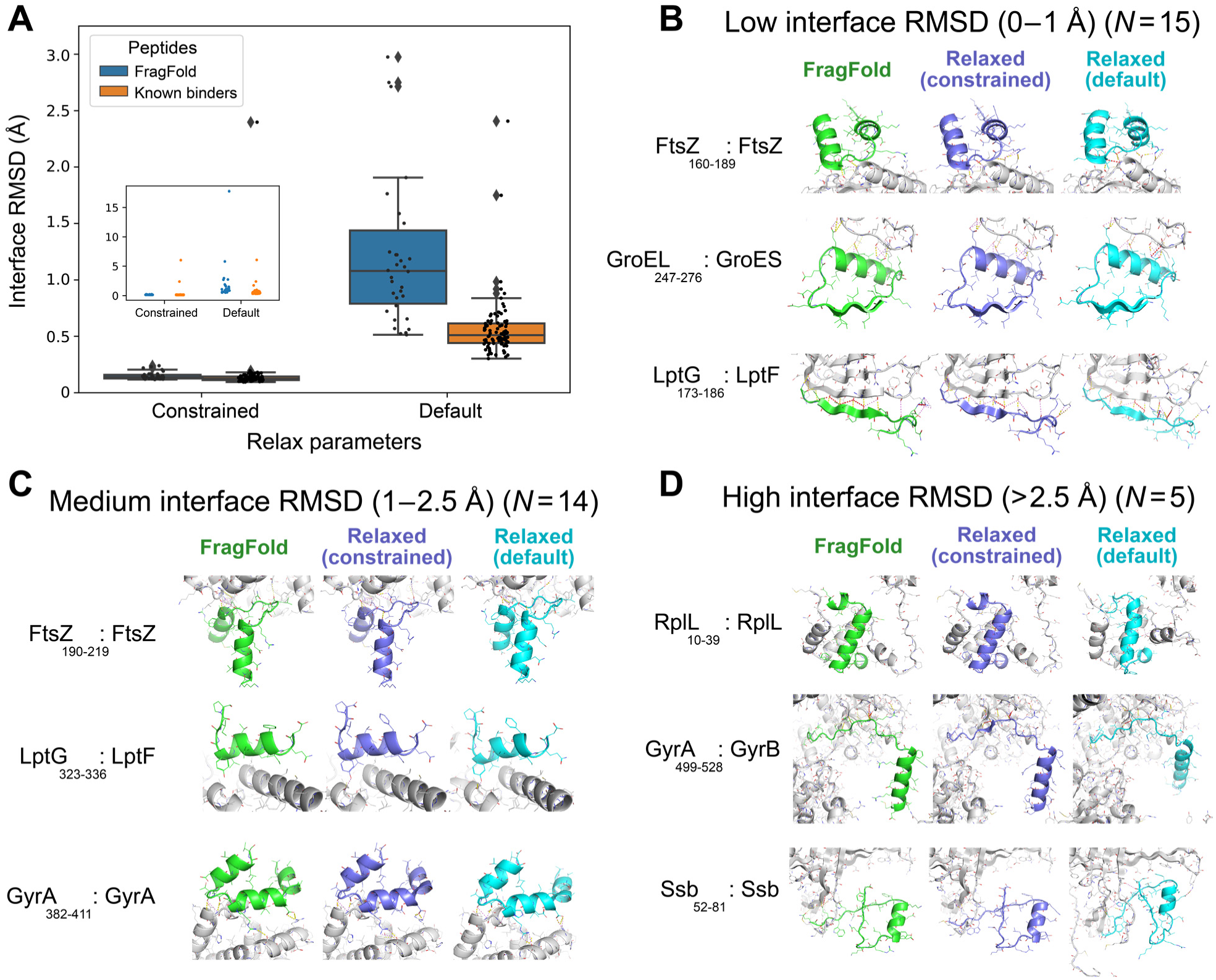
Rearrangements from Rosetta relaxation are typically minor for both FragFold models and experimental structures of protein-peptide complexes. (**A**) Box-and-whiskers plots of interface RMSD between the initial structure and the Rosetta forcefield-relaxed structure, for FragFold-predicted structures and experimentally determined structures from the PepPro database (45), for: *Left*: Rosetta relaxation with tight constraints; *Right*: Rosetta relaxation with default parameters. Inset: same plots, but also including 4 outliers with higher interface RMSD values. (**B**-**D**) Example FragFold structural models with no relaxation (fragment green), constrained Rosetta relaxation (fragment indigo), and default Rosetta relaxation (fragment cyan), for fragment-protein complexes exhibiting 0-1 Å (**B**), 1-2.5 Å (**C**), and >2.5 Å (**D**) interface RMSD between the FragFold structure and the (default parameters) Rosetta-relaxed model.

**Figure S9.**
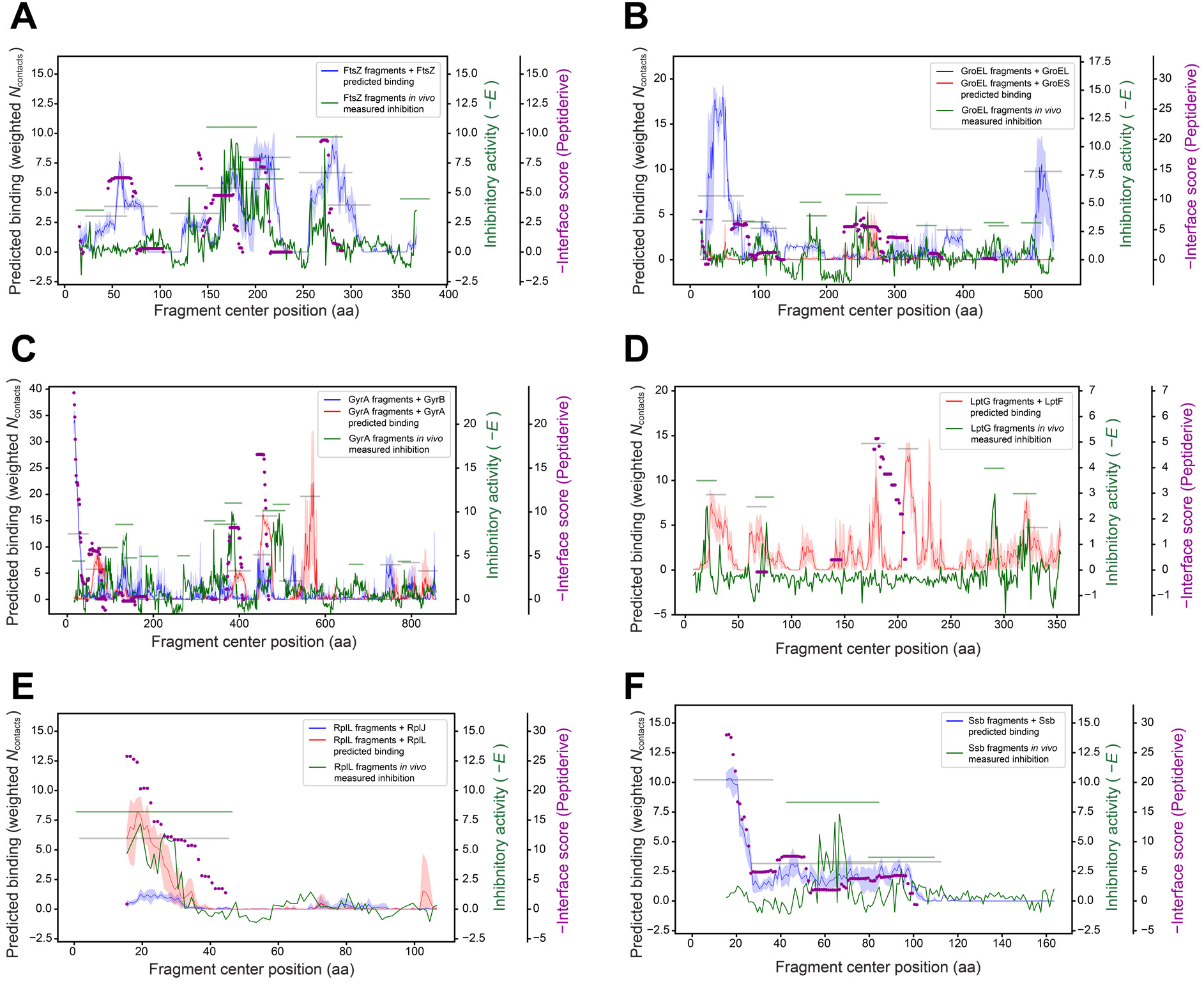
Peptiderive can predict some, but not all, inhibitory fragment peaks called by FragFold. (**A**) – (**F**) Plots of experimental inhibitory effects (green) and predicted binding from FragFold (blue and red), and automatedly called experimental and predicted peaks (green and grey bars), plotted as in Figure S2, overlaid with the predicted interface binding metric (-Interface score) from Peptiderive (purple dots); for fragments tiling across (**A**) FtsZ; (**B**) GroEL; (**C**) GyrA; (**D**) LptG; (**E**) RplL; and (**F**) Ssb, with a 1aa step size. Gaps in the Peptiderive data reflect that scores are only assigned to some regions of the sequence.

**Figure S10.**
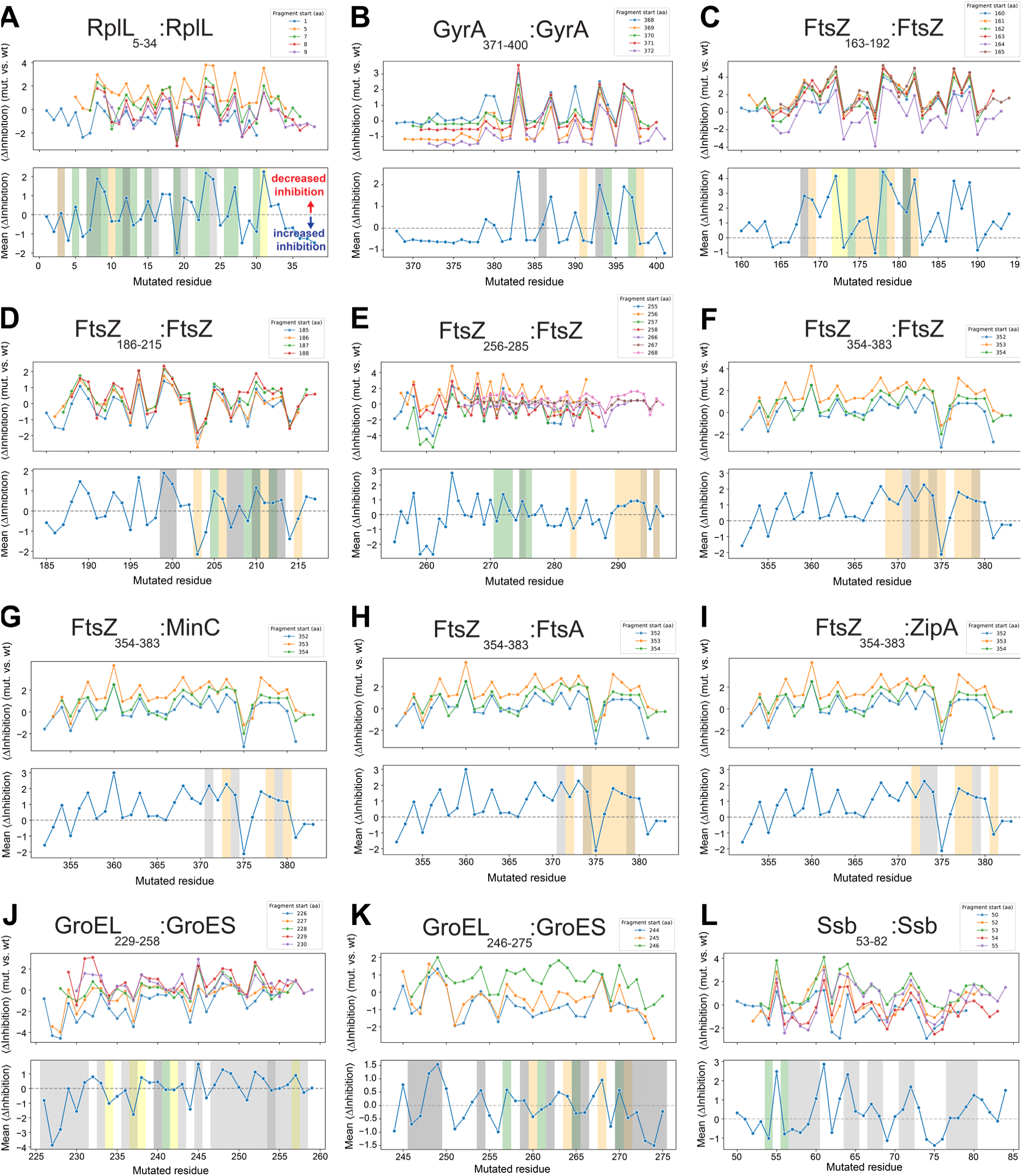
Mutational sensitivity plots for inhibitory fragment peaks from deep mutational scanning experiments. (**A**) – (**L**) Plots of mutational sensitivity (Δ_Inhibition_ = *E*_mutant_ – *E*_wt_)) for each inhibitory fragment peak corresponding to Fig. 6 (A) – (L), for both individual fragments spanning each peak (*upper*) and the mean across all peak fragments (*lower*). The mutational sensitivities measured for individual fragments tiling across each peak along with the average effect for each peak are shown. Contact residues are indicated with different shading colors overlaid on the plots of mutational sensitivity. Shading colors are partially transparent to allow overlap of residues that belong to multiple classes. **Yellow shading**: native full-length protein-protein interaction residues from experimental structures; **Orange shading**: FragFold model fragment-protein interaction residues; **Green shading**: interaction residues seen in *both* native protein-protein interaction structures and FragFold models of fragment-protein interactions; **Grey shading**: fragment *intramolecular* contact residues in either native full-length or FragFold-predicted structures; dark grey indicates fragment intramolecular contacts in *both* native and predicted structures.

### Supplementary Tables

**Table S1.** FragFold systematic predictions outperform a random model.

| **Fragmented protein** | $f_{\text{inhibitory peaks predicted}}$ | $f_{\text{predicted binding clusters inhibitory}}$ | $f_{\text{null, inhibitory peaks predicted}}$ | $f_{\text{null, predicted binding clusters inhibitory}}$ |
| --- | --- | --- | --- | --- |
| **All proteins** | **64%** | **68%** | **49%** | **51%** |
| **FtsZ** | 86% | 71% | 69% | 66% |
| **GroEL** | 44% | 67% | 39% | 55% |
| **GyrA** | 54% | 70% | 45% | 54% |
| **LptG** | 75% | 57% | 51% | 34% |
| **RplL** | 100% | 100% | 34% | 34% |
| **Ssb** | 100% | 67% | 63% | 42% |

**Table S2.** Systematic analysis shows FragFold predictions exhibit good specificity.

| Fragmented protein | Prediction type | $\rho_{\text{peaks}}$<br>(peaks/fragment/aa) | $\rho_{\text{on-target}} / \rho_{\text{off-target}}$<br>(specificity) | Partners |
| --- | --- | --- | --- | --- |
| FtsZ | on-target | 5.2E-05 |  | FtsZ |
| GroEL | on-target | 1.8E-05 |  | GroEL |
| GyrA | on-target | 7.1E-06 |  | GyrA, GyrB |
| LptG | on-target | 3.9E-05 |  | LptF |
| RplL | on-target | 3.8E-05 |  | RplL, RplJ |
| Ssb | on-target | 1.1E-04 |  | Ssb |
| **FtsZ** | off-target | 5.1E-06 | **10.0** | GroEL, GyrA, GyrB, LptF, RplL, RplJ, Ssb, eGFP |
| **GyrA** | off-target | 5.9E-07 | **11.9** | FtsZ, GroEL, LptF, RplL, RplJ, Ssb, eGFP |
| **RplL** | off-target | 9.6E-06 | **4.0** | FtsZ, GroEL, GyrA, GyrB, LptF, Ssb, eGFP |
| eGFP | control | 5.6E-06 |  | eGFP, GroEL, FtsZ, GyrA, GyrB, LptF, Ssb |

**Table S3.** Correlations between mutational sensitivity from deep mutational scanning experiments and structural models of fragment-protein complexes.

| Inhibitory peak | Structural model | TPR | TNR | PPV | NPV | MCC | *p* (MCC) |
| --- | --- | --- | --- | --- | --- | --- | --- |
| RplL<sub>5-34</sub> | FragFold | 0.54 | 0.60 | 0.41 | 0.71 | 0.13 | 0.12 |
| GyrA<sub>371-400</sub> | FragFold | 0.43 | 0.89 | 0.50 | 0.86 | 0.34 | 0.085 |
| FtsZ<sub>163-192</sub> | FragFold | 0.36 | 0.85 | 0.80 | 0.44 | 0.22 | 0.039 |
| FtsZ<sub>186-215</sub> | FragFold | 0.50 | 0.67 | 0.46 | 0.70 | 0.16 | 0.28 |
| FtsZ<sub>256-285</sub> | FragFold | 0.53 | 0.86 | 0.67 | 0.77 | 0.41 | 0.0012 |
| FtsZ<sub>354-383</sub> (+FtsZ) | FragFold | 0.50 | 0.93 | 0.90 | 0.59 | 0.46 | 0.012 |
| FtsZ<sub>353-382</sub> (+MinC) | FragFold | 0.33 | 1.00 | 1.00 | 0.54 | 0.42 | 0.021 |
| FtsZ<sub>354-383</sub> (+FtsA) | FragFold | 0.33 | 0.86 | 0.75 | 0.50 | 0.22 | 0.046 |
| FtsZ<sub>354-383</sub> (+ZipA) | FragFold | 0.33 | 0.93 | 0.86 | 0.52 | 0.31 | 0.088 |
| GroEL<sub>229-258</sub> | FragFold | 0.38 | 0.58 | 0.21 | 0.75 | -0.04 | 0.74 |
| GroEL<sub>246-275</sub> | FragFold | 0.75 | 0.42 | 0.30 | 0.83 | 0.15 | 0.098 |
| Ssb<sub>53-82</sub> | FragFold | 0.25 | 0.74 | 0.33 | 0.65 | -0.01 | 0.68 |
| RplL<sub>5-34</sub> | Native | 0.62 | 0.64 | 0.47 | 0.76 | 0.24 | 0.032 |
| GyrA<sub>371-400</sub> | Native | 0.43 | 0.96 | 0.75 | 0.87 | 0.49 | 6.6 x 10<sup>-4</sup> |
| FtsZ<sub>163-192</sub> | Native | 0.18 | 0.85 | 0.67 | 0.38 | 0.04 | 0.26 |
| FtsZ<sub>186-215</sub> | Native | 0.42 | 0.81 | 0.56 | 0.71 | 0.24 | 0.037 |
| FtsZ<sub>256-285</sub> | Native | 0.13 | 0.89 | 0.40 | 0.66 | 0.04 | 0.22 |
| GroEL<sub>229-258</sub> | Native | 0.62 | 0.50 | 0.28 | 0.81 | 0.11 | 0.15 |
| GroEL<sub>246-275</sub> | Native | 0.62 | 0.50 | 0.29 | 0.80 | 0.11 | 0.16 |
| Ssb<sub>53-82</sub> | Native | 0.25 | 0.74 | 0.33 | 0.65 | -0.01 | 0.67 |

Metrics describing the correlations between (1) mutationally-sensitive residues with $\langle \Delta_{\text{Inhibition}} \rangle > 0.5$ , corresponding to sites where mutations decreased inhibitory activity, from experimental deep mutational scanning data on inhibitory protein fragments; and (2) residues making fragment:protein contacts and fragment intramolecular folding contacts in structural models of fragment:protein complexes. Structural models tested included FragFold models (top set of entries) and experimental structures of native full-length protein-protein complexes (bottom set of entries). TPR, true positive rate (fraction sensitive residues that were binding residues in model); TNR, true negative rate (fraction non-sensitive residues that were non-binding residues in model); PPV, positive predictive value (number of true positives / (number of true positives + number of false positives)); NPV, negative predictive value (number of true negatives / (number of true negatives + number of false negatives)); MCC, Matthews correlation coefficient between the structural model and the experimental mutational sensitivity.

## Notes

https://github.com/swanss/FragFold

https://figshare.com/articles/dataset/Source_Data_for_Savinov_and_Swanson_et_al_2023/24841269

### Competing Interest Statement

The authors have declared no competing interest.

### Summary of Updates

The revised manuscript is expanded with many new results, arising from additional computational analyses and new experimental data. A brief summary of the major additions follows: 1. We now include extensive experimental deep mutational scanning data on several inhibitory protein fragments. These data reveal the accessibility of mutations yielding further improved inhibitory protein fragments, with implications for designing peptide-based therapeutics; and provide systematic experimental support for the FragFold-predicted fragment binding modes, including for the novel binding modes predicted for the C-terminal fragment of FtsZ. 2. We perform an analysis of predicted fragment binding energy with Rosetta to show that the predicted fragment binding modes are physically plausible and broadly consistent with those seen for known peptide binders with experimentally-determined structures. Furthermore, relaxation of fragment binding modes in the Rosetta force field does not substantially change binding conformations or binding energy, suggesting the binding modes directly predicted by AlphaFold are reasonable without fine-tuning. 3. We perform additional extensive FragFold predictions against non-cognate protein partners to further demonstrate prediction specificity. 4. We compare the performance of FragFold with an alternative approach, the Rosetta-based Peptiderive protocol. 5. We have included numerous additional changes in response to a public review of the original bioRxiv manuscript (Version 1) from CJ San Felipe, Priyanka Bajaj, and James Fraser. These include clarifications of how MSAs and sequence inputs were treated in FragFold predictions.

